# Osmotic Stress Influences Microtubule Drug Response Via WNK1 Kinase Signaling

**DOI:** 10.1101/2024.07.08.602030

**Authors:** Ana Monfort-Vengut, Natalia Sanz-Gómez, Sandra Ballesteros-Sánchez, Beatriz Ortigosa, Aitana Cambón, Maria Ramos, Ángela Montes-San Lorenzo, Juan Manuel Rosa-Rosa, Joaquín Martínez-López, Ricardo Sánchez-Prieto, Rocío Sotillo, Guillermo de Cárcer

## Abstract

Ion homeostasis is critical for numerous cellular processes, and disturbances in ionic balance underlie diverse pathological conditions, including cancer progression. Targeting ion homeostasis is even considered as a strategy to treat cancer. However, very little is known about how ion homeostasis may influence anticancer drug response. In a genome-wide CRISPR-Cas9 resistance drug screen, we identified and validated the master osmostress regulator WNK1 kinase as a modulator of the response to the mitotic drug rigosertib. Osmotic stress and WNK1 inactivation lead to an altered response not only to rigosertib treatment but also to other microtubule-related drugs, minimizing the prototypical mitotic arrest produced by these drugs. This effect is due to an alteration in microtubule stability and polymerization dynamics, likely maintained by fluctuations in intracellular molecular crowding upon WNK1 inactivation. This promotes resistance to microtubule depolymerizing drugs, and increased sensitivity to microtubule stabilizing drugs. In summary, our data proposes WNK1 osmoregulation activity as a biomarker for microtubule-associated chemotherapy response.

## 1. INTRODUCTION

Targeting mitosis is one of the most efficient strategies to stop cancer cell proliferation (1–3), and in the last decades a wide variety of new mitotic drugs were developed targeting essential regulator proteins that govern mitosis progression (polo-like kinases, aurora kinases, nek kinases, kinesins, etc.) (4–6). Despite many of these new mitotic drugs are very efficient in killing cancer cells in vitro, the results from the clinical trials were not successful and none of these compounds are approved for cancer therapy (7, 8). A perfect example of this paradigm is the drug rigosertib (ON-01019). It was originally described as a Plk1 kinase inhibitor, but in recent years it has become clear that rigosertib is a multi-target drug, inhibiting multiple pathways from PI3K-AKT to RAS-MEK-ERK signaling, or cellular processes such as microtubule dynamics, with a strong controversy on the precise mechanism of action (9). Despite its efficacy in preclinical studies and the low cytotoxicity shown in clinical trials, rigosertib only showed moderate results in targeting myelodysplastic tumors not reaching the clinical setting (10). The fact that the exact molecular mechanism is not known, as well as the absence of appropriate biomarkers, may explain the lack of success of rigosertib in the clinic. To define rigosertib biomarkers, we performed a resistance genome-wide CRISPR-Cas9 screen, in breast cancer cell lines, and identified the downregulation of the “With-No-lysine [K] kinase 1” (WNK1) as the mayor hit for rigosertib resistance.

WNK1 is a master regulator of ion homeostasis, it controls the influx/efflux of chloride ions upon osmotic stress induction, and it is related to blood pressure control diseases (11–14). Chloride ions (Cl^-^) are responsible for osmotic pressure and acid-base balance, which determine fundamental biological functions in all tissues (15), and impaired Cl^-^ balance affects several essential cellular processes underlying many pathological conditions, from neuronal excitability to heart failure, cystic fibrosis, or chronic kidney disease (16). Regarding cancer, ion homeostasis is involved in the proliferation, apoptosis, migration or angiogenesis of cancer cells (17, 18). Importantly, WNK1 is also related to cancer progression (19), and is even considered a putative therapeutic target (20, 21). Although ion homeostasis is also described to influence chemoresistance (22, 23), little is known about the role of WNK1 in cancer therapy resistance.

Here we describe how WNK1 inactivation leads to rigosertib resistance in breast cancer cell lines, due to alteration in microtubule polymerization dynamics, confirming that rigosertib mainly acts as a microtubule poison. Furthermore, WNK1 downregulation alters the cellular response to general microtubule-related drugs. We propose that WNK1 inactivation, and the subsequent osmotic stress, lead to changes in intracellular molecular crowding that alter microtubule polymerization dynamics, conferring resistance to microtubule depolymerizers and sensitivity to microtubule stabilizers.

## 2. MATERIAL AND METHODS

### 2.1. - Mammalian cell line culture

Cell lines were sourced from the American Type Culture Collection (ATCC®). MDA-MB-453 (RRID:CVCL_0418), MCF-7 (RRID: CVCL_0031), EVSA-T (RRID: CVCL_1207), MDA-MB-231 (RRID: CVCL_0062), HEK-293 (RRID:CVCL_0063), and RPE-1 (RRID:CVCL_4388) cell lines were grown adherently in Dulbecco’s Modified Eagle Medium (DMEM) supplemented with 10% fetal bovine serum (FBS), in a 37°C and 5% CO2 atmosphere, and routinely tested for the presence of Mycoplasma species. Cell line authentication was done with the GenePrint® 10 System (Promega), and data were analyzed using GeneMapper® ID-X v1.2 software (Applied Biosystems).

Osmotic stress was induced by either using regular DMEM media for the isoosmotic condition (∼335 mOsm/kg); DMEM diluted with 1/3 of milliQ water for the hypoosmotic condition (∼225 mOsm/kg) (24); or DMEM supplemented with 1.5% (wt/vol) of 300 Da polyethylene glycol (PEG300) (VWR, VWRC26605.290) for the hyperosmotic condition (∼415 mOsm/kg) (25). In all cases, 10% FBS was added once media were prepared to not interfere with the final growth factor concentration. Media osmolarity was confirmed using a freezing-point osmometer (Gonotec®, Osmomat 010).

### 2.2.– Lentiviral particle generation and cell infection

Third generation lentiviral particles were produced by transfection of HEK-293 cells with a mixture of 10 μg of the lentiviral plasmid of interest, and 6.5 μg, 2.5 μg, and 3.5 μg of the lentiviral packaging vectors (pMDL, pRev, and pVSVg respectively) per p100 plate, using Lipofectamine 2000 (ThermoFisher, 11668019). Cell media was changed 6 h after transfection and cells were incubated at 37° C to allow for virus production. Media containing lentiviral particles was harvested at 48 h and 72 h, filtered through 0.45 μM PVDF membrane filters and flash-frozen at −80° C. Cell infection was performed by adding lentiviral particles to cell lines in the presence of 8 μg/ml polybrene (Sigma, TR-1003) for 12 h followed by fresh media addition. The selection was further done in the presence of antibiotics such as hygromycin B (50 μg/ml), G418 (400 μg/ml), or puromycin (2 μg/ml), depending on the selection marker.

### 2.3.– CRISPR-Cas9 genome-wide library generation

TET-ON-Cas9-inducible expressing cells were generated firstly by infection with the rtTA plasmid pLVX-Tet3G (Takara, 631358) and neomycin selection, followed by infection with the U6-sgRNA-TRE-Cas9-P2A-dsRED-EFSGFP/Puro lentiviral particle (LC-TRIP) (26), and puromycin selection, and single-cell cloning to ensure Cas9 expression homogeneity. The sgRNA library, in the pKLV2-U6gRNA5(BbsI)-PGKpuro2ABFP-W plasmid Addgene (#67989) (27), was amplified following the detailed protocol described by Koike-Yusa and colleagues (28), and titrated using BFP detection by flow cytometry (29). The gRNA library was then transduced in the MDA-MB-453 Cas9-expressing cell line by infecting 30×10^6^ cells (100x sgRNA coverage) with a 0.3 multiplicity of infection (MOI) plus polybrene (8 μg/ml) during 2 h. BFP-positive cells were sorted and frozen always keeping a minimum of 10×10^6^ cells per cryovial to keep a minimum of 50-fold sgRNA coverage.

### 2.4.– Rigosertib drug resistance screen

A minimum of 5×10^6^ cells (to ensure 50-fold library coverage) of the MDA-MB-4563 cell line CRISPR library were treated with doxycycline (1 μg/ml) for 7 days to induce Cas9 expression and library genetic edition. Cas9 active cells were then treated with rigosertib (100 nM) for 3 days followed by a 4-day holiday period. This protocol was repeated five times until resistant colonies appeared. Once clonal-resistant colonies appeared, 24 resistant colonies were individually isolated and expanded for further analysis. In addition, the rest of the positive colonies were pooled and genomic DNA was extracted for sgRNA enrichment identification

### 2.5.– Drug screen sgRNA identification

sgRNA identification in the isolated rigosertib-resistant colonies was done by PCR amplification of the pKLV2 plasmid sgRNA insert region, and PCR product subcloning in the pGEM-T Easy Vector Systems (Promega #A1360) followed by sanger sequencing of pGEM-T positive bacterial colonies. The sgRNA identification, from the pooled resistant cells, was done by PCR amplification of the pKLV2 plasmid sgRNA insert region, using the Q5 High-Fidelity DNA polymerase (NEB, M0491), and the following primers/barcodes (table 1):

**Table 1.**
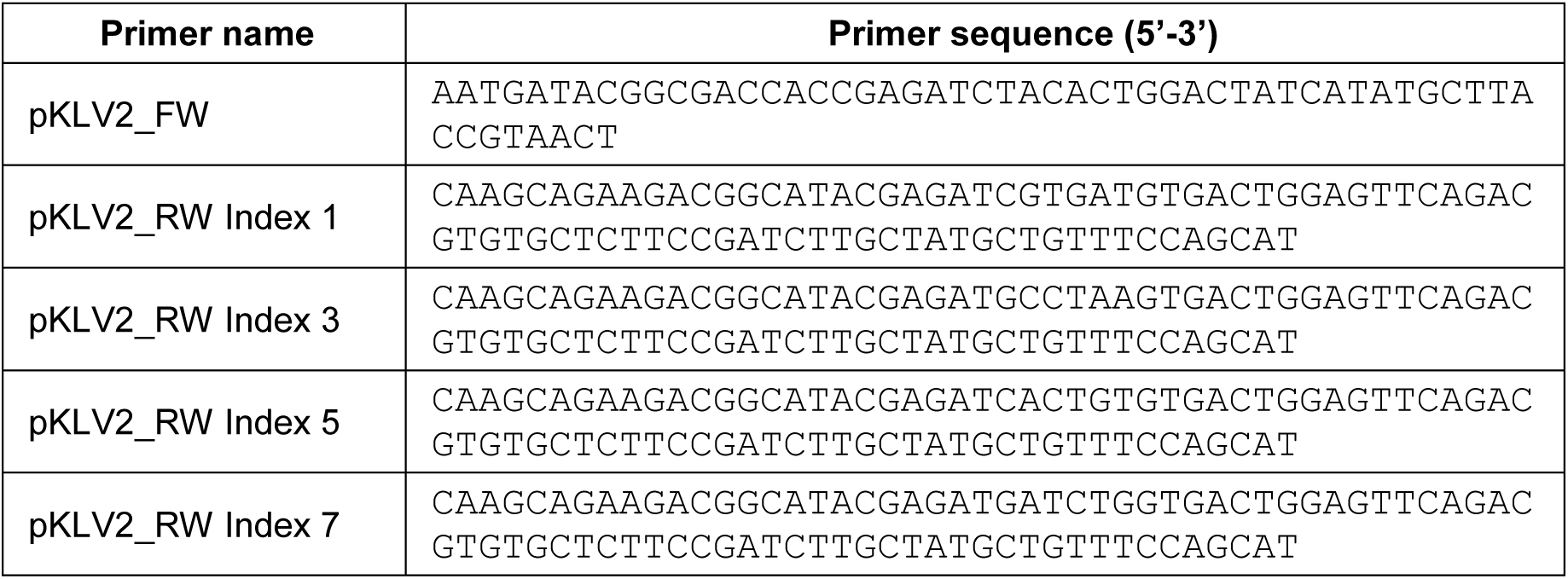
PCR primers with barcodes/indexes used for sgRNA IonTorrent sequencing.

Amplified DNA was purified using AMPure XP Bead-Based Reagent (Beckman Coulter, A63880). For the sequencing library preparation, PCR samples were treated with NEBNext End Repair Module (NEB, E6050) and the adaptor ligated DNA was amplified using NEBNext Q5 Hot Start Hifi PCR Master Mix (NEB, M0543). The libraries were sequenced at a rate of 5 million reads per sample, using the Ion Torrent PGM Platform and the Ion 540 Ampliseq Chip kit (ThermoFisher, A27765).

### 2.6.– WNK1 genetic depletion

WNK1 depletion by CRISPR-Cas9 editing was done by using two independent gRNAs targeting Exon-1 (5’-CGCCGACGCTGTGACCGGC-3’) and Exon-4 (5’-ACTTACACTGGTCACGCGA-3’) cloned into pKLV2-U6gRNA5(BbsI)-PGKpuro2ABFP-W backbone vector (Addgene #67974). TET-ON-Cas9 expressing cells were infected and BFP-positive cell FACS sorted. WNK1 depletion was done by incubating cells with doxycycline (1 μg/ml) for 7 days. To avoid possible overgrowth of WNK1-WT escapers, WNK1-depleted cells were transiently generated before each experiment. WNK1 edition efficiency was confirmed using the T7-Endonuclease I assay, by PCR amplification of the specific sgRNA target region in the WNK1 locus, exon 1 (FW 5’-GGCTTGTCGGTGCTGAGTG-3’ and REV 5’-CTCCCCACAAGGCTAGGCT-3’) and exon 4 (FW 5’-GCCCCTGAGATGTATGAGGAG-3’ and REV 5’-CCCAACCCCAACAGTTCAC-3’). PCR fragments were incubated with the T7 Endonuclease I (NEB, M0302S) and nuclease activity was confirmed by DNA agarose electrophoresis.

WNK1 genetic indels were identified by pGEM-T subcloning of the same PCR products generated for the T7-endonuclease assay, and positive bacterial colonies were subjected to plasmid DNA sequencing.

WNK1 mRNA silencing was done by using shRNAs lentiviral pLKO plasmids (SIGMA) TRCN0000219718 (5’-TCTGCGGGAAGGCGGTTTATA-3’) and TRCN0000000919 (5’-CCGCGATCTTAAATGTGACAA-3’) and puromycin selection for 72 h.

### 2.7.– RNA-seq and qRT-PCR

RNA from control cells or the rigosertib-resistant colonies was extracted with the mirVana miRNA extraction kit (AM1571, Invitrogen) following the manual instructions, and 250 ng with an average RIN=9.6 (range: 9.4-9.8) in a labchip analysis, were processed into cDNA sequencing libraries with the “QuantSeq 3‘ mRNA-Seq Library Prep Kit (FWD) for Illumina” (Lexogen, Cat.No. 015). Single-stranded sequencing was carried out with an Illumina NextSeq 500 sequencer obtaining about 10 million reads per sample. Reads were analyzed for quality with FastQC and aligned against the NCBI reference human genome build GRCh38.p13 with HiSat2 (30). Reads were identified with the GTF coordinate file and counted with HTSeq software (31). Raw counts were normalized to median ratios in the DeSeq2 package (32). Differential expression analysis was performed with DeSeq2 comparing rigosertib resistant colonies to control samples, and the volcano plot was generated with the EnhanceVolcano software package. RNA-seq data have been deposited at the Gene Expression Omnibus (GSE271314). For qRT-PCR, RNA extraction was done using the RNeasy Micro Kit (Qiagen, 74004). cDNA synthesis was done using the QuantiTect Reverse Transcription Kit (Qiagen, 205311). Real-time PCR was done on a starting material of 15 ng of cDNA with SYBR Green PCR Master Mix (Applied Biosystems, 4309155) in a LightCycler II® 480 (Roche), using the WNK1 primers (FW: 5’-CAGAGCCCTGGAATGAACTTG-3’, and REV 5’-TTAGGAGGGCTGCTTGTTGC-3’) and actin primers(ACTB) as housekeeping gene (FW 5’-TGGATCAGCAAGCAGGAGTATG-3’ and REV 5’-GCATTTGCGGTGGACGAT-3’).

### 2.8.– Small Compounds

Small compounds, listed in table 2, were used at the doses indicated in figure legends and main text. All compounds were dissolved in DMSO, with the exception of doxycycline that was dissolved in sterile water, and stored at −20° C. To avoid rigosertib degradation, powder aliquots (0.5 mg each) were dark-stored in opaque tubes and conserved at −80°C. A 1 mM master stock was freshly done for every experimental setting.

**Table 2:** drug name, vendor source and reference number.

| Drug name | Source | Reference |
| --- | --- | --- |
| ABT-751 | MedChemExpress | HY-13270 |
| Colchicine | Sigma Aldrich | C9754 |
| Danuserib | Selleckchem | S1107 |
| DIDS-sodium salt | MedChemExpress | HY-D0086 |
| Doxorubicin | Sigma Aldrich | D1515 |
| Doxycycline | BioChemica | A2951 |
| Nocodazole | Selleckchem | S2775 |
| Paclitaxel | Selleckchem | S1150 |
| Rapamycin | LC Laboratories | R-5000 |
| Rigosertib | MedChemExpress | HY-12037A |
| Torin1 | Selleckchem | S2827 |
| Trametinib | Selleckchem | S2673 |
| Volasertib | Selleckchem | S2235 |
| WNK-463 | MedChemExpress | HY-100626 |
| WNK-IN-11 | MedChemExpress | HY-112094 |

### 2.9.– Cell viability assays

Cell growth curves were done by seeding cells in 6-well plates (3×10^5^ cells/well) in triplicates and counting every three days using the 3T3 protocol (1/3 dilution every 3 days).

Colony formation assays were done by seeding in 12-well plates (2×10^3^ cells/well), or in 24-well plates (1×10^3^ cells/well). Drugs were added at the indicated concentrations and renewed every 3 to 4 days. After 10 to 14 days, or until the control wells (DMSO) were 80-100% confluent, cells were fixed with 4% formaldehyde in PBS, stained with Giemsa, and plates scanned in an Epson V800 scanner. The colony area was quantified using Image J (NIH Bethseda, MD) and its ColonyArea plugin (33). IC50 plots were calculated using the Prism software (GraphPad, Boston, MA)

For the competition assays, cells carrying inducible Cas9 and the WNK1 sgRNAs were infected with either EGFP-expressing lentiviral particles (FUGW-EGFP, Addgene #14883) or FUGW-mRuby3-expressing lentiviral particles (34). EGFP-positive cells were doxycycline treated for 7 days, to deplete the WNK1 gene (WNK1-KD), and then mixed with mRuby3 cells with unedited WNK1 gene (WNK1-WT) at a proportion of 20%-80%. 3×10^5^ of the cell mixture was seeded in p6 wells by triplicates and incubated in the presence of DMSO or the indicated drugs. Cells were grown for 18-29 days with drug renewal every 3 to 4 days. EGFP vs. mRuby3 proportions were measured by flow cytometry CytoFLEXTM (Beckman Coulter) every 72-96 h by cell suspension in sorting buffer (PBS, 2 % BSA, 1 mM EDTA, 1 mM HEPES pH 7.4).

### 2.10.– Cell cycle profiling

Cell cycle profiling was done by measuring DNA content in the assayed cells by flow cytometry. Trypsinized cells were fixed in slow agitation with cold 70° ethanol and kept on ice for 30 minutes. Then, washed twice with PBS-Triton X-100 0.03% (Fisher BioReagents, 10102913), and DNA was counterstained with DAPI (1 μg/ml) (AppliChem, A4099) in PBS-Triton X-100 0.03%. DNA signal was estimated by fluorophore excitation with a 405 nm violet laser and the 450/50 filter in a BD FACSCanto II Flow Cytometer (BD).

### 2.11. Immunofluorescence and microtubule polymerization assays

Cells growing on coverslips were fixed in 4% methanol-free formaldehyde (PolySciences, 18814) in PBS and permeabilized with cold methanol. Then, they were blocked with 10% FBS in PBS-T (PBS, triton-X100 0.03%) and incubated for 2 h with the primary antibodies, described in table 3, diluted in PBS-T. Then secondary antibodies, coupled with Alexa488 (green) or Alexa546 (red) dyes (ThermoFisher), were diluted in PBS-T and incubated for 1 h. Finally, DNA was counterstained with DAPI (0.1 μg/ml), and cells were mounted in glass slides using ProLong Diamond antifade mounting media (ThermoFisher, P36965). Pictures were obtained using a Nikon ECLIPSE 90i microscope. For detailed mitotic fate evaluation, pictures were taken with a LSM710 (Zeiss) confocal laser scanning microscope.

**Table 3:** Antibodies used in different applications: WB: Western blot / IF: Immunofluorescence / IHC: immunohistochemistry

| Antibody (clone) | Source | Reference | Host species | Application | Dilution |
| --- | --- | --- | --- | --- | --- |
| alpha-tubulin (DM1A) | Santa Cruz | sc-32293 | Mouse | IF / WB | 1:2000 |
| gamma-tubulin (GTU88) | Sigma Aldrich | T6557 | Mouse | IF | 1:1000 |
| acetylated-alpha-tubulin | Sigma Aldrich | T7451 | Mouse | WB | 1:500 |
| detyrosinated-alpha-tubulin | Sigma Aldrich | AB3201 | Rabbit | WB | 1:500 |
| histone H3 phospho-S10 | Sigma Aldrich | AB3201 | Rabbit | IF | 1:1000 |
| MDR1 | Cell Signaling | #12683 | Rabbit | WB | 1:500 |
| SPAK | MRC PPU | 588713 | Sheep | WB | 1:2000 |
| SPAK phospho-S373 | MRC PPU | 588719 | Sheep | WB | 1:500 |
| WNK1 | ThermoFisher | PA5-28382 | Rabbit | WB | 1:500 |
| ERK5 phospho-T218/Y220 | Cell Signaling | #3371 | Rabbit | WB | 1:1000 |
| GAPDH (FF26A) | CNIO | - | Mouse | WB | 1:10000 |
| vinculin (7F9) | Santa Cruz | sc-73614 | Mouse | WB | 1:2000 |
| histone H3 phospho-S10 | Merck | 06-570 | Rabbit | IHC | 1:200 |
| C3 Cleaved Caspase-3 | Cell Signaling | # 9661 | Rabbit | IHC | 1:300 |

Microtubule polymerization assays were done in RPE-1 cells incubated with the WNK1 inhibitor WNK-463 (2 μM) or DMSO (control) for 24h. Cells were then chilled over an ice water bath for 30 min, to fully depolymerize the microtubule lattice, and then brought back into warm (37° C) DMEM for 1 min, allowing microtubule regrowth. Immediately, cells were treated for 3 min in CSK buffer (10 mM PIPES/KOH pH 6.8, 100 mM NaCl, 300 mM Sucrose, 1 mM EGTA, 1 mM MgCl2, 1 mM DTT) supplemented with Triton-X100 0.03% to extract the soluble depolymerized tubulin. Then cells were fixed in absolute cold methanol and stored at −20° C. As a control for the depolymerization assay, some coverslips skipped the ice treatment and were exclusively treated with CSK+Tx 0.03% buffer and fixed. Cells were then subjected to alpha- and gamma-tubulin immunofluorescence and microtubule regrowth capacity was quantified by measuring the alpha-tubulin intensity signal in a 20 μm^2^ rounded ROI centered at the gamma-tubulin signal, in about 50 to 100 cells per condition, using ImageJ software program (NIH, Bethseda, MD). The same procedure was followed for WNK1-WT and WNK1-KD MCF-7 cells.

### 2.12. Protein lysis and Immunoblotting

For MDR1 and WNK1 detection, cells were lysed with Laemmli buffer (2% SDS, 10% Glycerol, 60 mM Tris pH 6.8), boiled at 100°C for 10 min, and centrifuged at 13300 rpm for 15 min. For the rest of proteins, cells were lysed in RIPA buffer (20 mM Tris-HCl pH 7.5, 150 mM NaCl, 1mM EDTA, 1 mM EGTA, 1% NP-40, 1% Sodium deoxycholate) supplemented with 50 U/mL Benzonase® Nuclease (Sigma, #E1014) and 1% proteases and phosphatases inhibitors (Merck, #P3840; Sigma, #P0044; Sigma, #P5726) on ice for 20 minutes, vortexing every 5 minutes, and centrifuged at 4° C for 15 min at 13300 rpm. Lysate supernatants were quantified for protein concentration using the BCA Protein Assay Kit (TermoFisher Scientific, 23225) and separated on 4-20% Tris-Glycine gels (NovexTM, Life Technologies). In the case of WNK1, SPAK and pSPAK-Ser373, gels were transferred onto PVDF membranes with a constant ampere of 250 mA for 16 h in transfer buffer (Tris/Glycine, 10% methanol and 0.1% SDS). For the other antibodies, proteins were transferred onto nitrocellulose membranes using Trans-Blot® TurboTM Transfer System RTA Transfer Kit (BioRad). Blotted membranes were blocked in 10% non-fat milk in PBS-Tween 20 0.05% for 30 minutes at room temperature, and then incubated at 4° C overnight with the primary antibody (table 3). The corresponding secondary antibodies, coupled to fluorescent IRDye680, were incubated for 45 minutes, and scanned with the Odyssey Infrared Imaging System (Li-Cor Biotechnology).

### 2.13. Mouse xenograft experiments

Mouse tumoral xenograft experiments were done with 8-10 week NMRI female nude mice acquired from Charles Rivers, and they were maintained under standard housing conditions with free access to chow diet and water, as recommended by FELASA (Federation of European Laboratory Animal Science Association). MDA-MB-453 WNK1-WT and WNK1-KD cells were tested for Mycoplasma before engrafting. Briefly, 1×10^6^ cells in 100 μl serum-free DMEM were injected in each mouse flank, MDA-MB-453 WNK1-WT in the left flanks and WNK1-KD in the right flanks. When the tumors reached approximately 100 mm^3^, they were randomly assigned to rigosertib or vehicle group (6 mice per group), and treatment started with rigosertib sodium salt (300 mg/kg) or the vehicle solution via oral gavage for 5 days followed by 2 days of holiday, during 3 consecutive weeks. Tumors were also measured daily using a caliper and tumor volume was calculated applying the formula: Volume = ½ (Length x Width^2^). At the end of the treatment, mice were euthanized, and tumors were formalin-fixed and embedded in paraffin. Mice work was performed under the ethical approval protocol G-18/21 from the DKFZ (Heidelberg, Germany) Animal Welfare Office.

To discard rigosertib toxicity, mice weight was measured daily once the drug was administered. In addition, weekly analysis of blood cell populations was performed by extracting fresh blood in K3-EDTA 3K (Aquisel, 1501126) sample tubes, and analyzed using a hemocytometer (Scil Vet abc Plus, Antech).

### 2.14. Tumor tissue immunohistochemistry

Mice tumoral tissue paraffin sections were subjected to deparaffinization with xylene and rehydration with graded ethanol. Antigen retrieval was done under 0.09% (v/v) unmasking solution (Vector Labs) for 30 minutes in a steamer, followed by inactivation of endogenous peroxidases using 3% hydrogen peroxide (Sigma) for 10 min. Primary antibodies used were phospho-histone H3-Ser10 and C3 cleaved Caspase 3. Specific Alexa fluorophore-labeled goat IgG was used as a secondary antibody, and DAB Peroxidase Substrate kit (Vector Labs) was utilized for antibody detection. Tumor sections were visualized under a Leica DMi1 inverted microscope and analysis of images was performed using ImageJ software program (NIH, Bethseda, MD).

### 2.15. Statistical analysis

Statistical analysis was performed with the Prism software (GraphPad) using different methods as described in the figure legends. Probabilities of less than 0.05 were considered statistically significant (*p < 0.05; **p < 0.01; ***p < 0.001; ****p < 0.0001).

## 3. RESULTS

### 3.1. CRISPR-Cas9 screening identifies WNK1 as a determinant for rigosertib response

The Yusa sgRNA library (27) was introduced in the breast cancer cell line MDA-MB-453 with inducible expression of Cas9. The sgRNA MDA-MB-453 library was assayed against 100nM of rigosertib for three days, and a holiday recovery time of four days, repeating this schedule five times until resistant colonies were visible (Figure 1A, and Supp. Figure 1A). A collection of 12 resistant colonies was isolated, and the rest of the resistant cells were pulled for further genomic analysis. Isolated resistant colonies were then corroborated to be refractory for rigosertib showing a 3 to 5-fold increase in the rigosertib IC50 compared to the parental cells (Figure 1B). None of the isolated rigosertib-resistant cells expressed the multi drug resistant protein (MDR1) (35) (Supp. Figure 1B), and they were as sensitive as the parental cells to other classical anticancer drugs such as the genotoxic agent doxorubicin, the Plk1 mitotic kinase inhibitor volasertib, or the MEK inhibitor trametinib (Supp. Figure 1C), indicating that cell survival was not based on a general drug resistance mechanism. We also confirmed the acquired resistance by checking the capacity of rigosertib to arrest the cell cycle in mitosis (36–38). Resistant colonies showed no cell cycle arrest in the 4N DNA content (Figure 1C), when compared to parental cells, displaying a significant reduction in mitotic arrest as depicted by histone H3 phospho-Ser10 (p-HH3) staining (Figure 1D).

**Figure 1.**
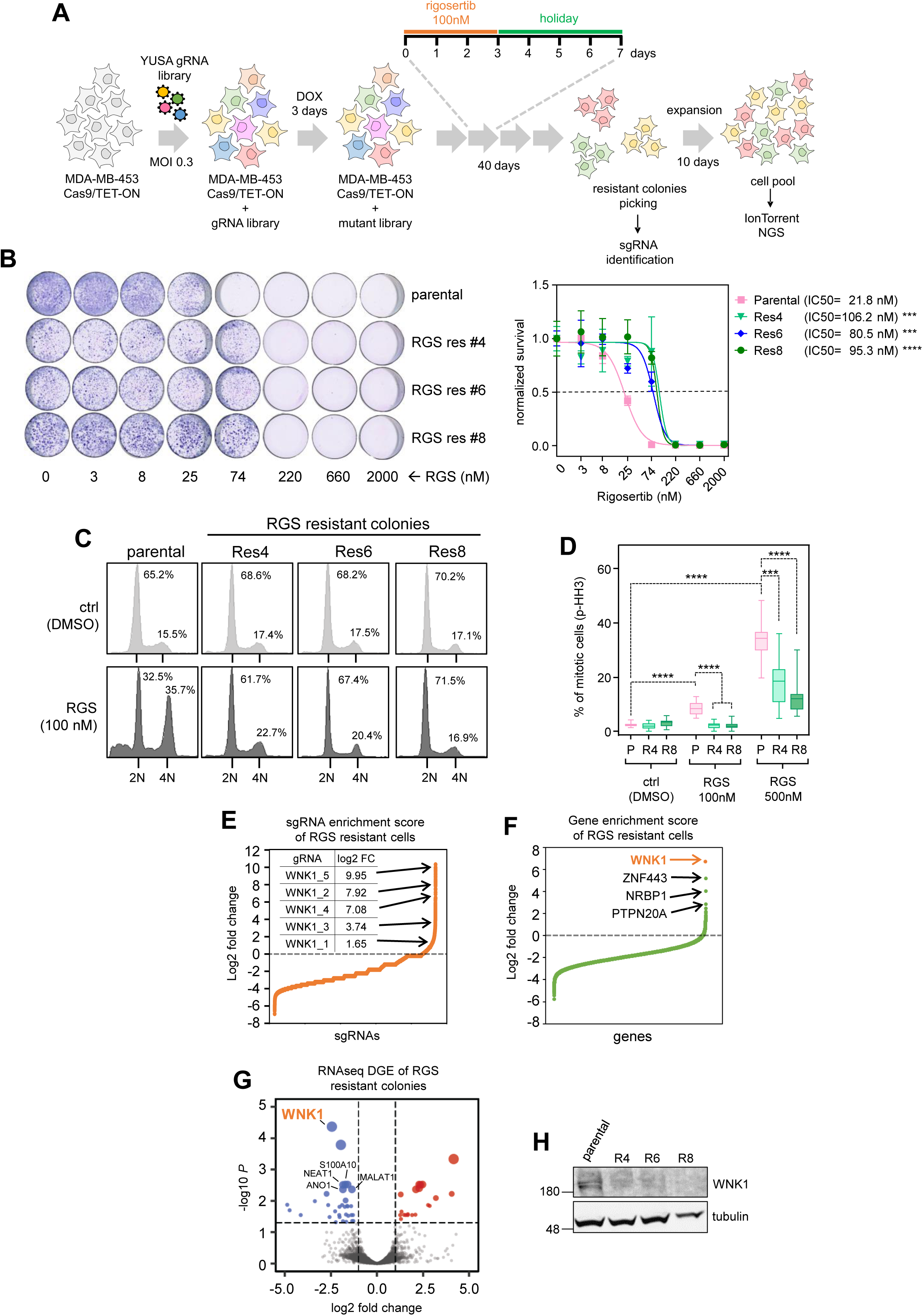
CRISPR-Cas9 screening identifies WNK1 as a hit for rigosertib-acquired resistance: **(A)** Cartoon summarizing the CRISPR-Cas9 screening process. Three days after Cas9 activation (+DOX), the sgRNA library was subjected to 100nM rigosertib for 3 days followed by 4 days of holiday. This drug regime was repeated five times allowing resistant colonies to appear. Then proliferating colonies were isolated, and the rest of the resistant cells were pooled. **(B)** Colony formation assays of the isolated rigosertib-resistant colonies (RGS res) in the presence of serial dilutions of rigosertib (left panel), and normalized logarithmic regression plot for rigosertib IC50 calculation of the parental MDA-MB-453 cells (pink line) and the resistant colonies (blue and green lines). Two-way ANOVA with Tukey multiple comparisons test of resistant versus the parental cells: *p*<0.001 (***) and *p*<0.0001 (****). Representative graph of three experimental replicates. **(C)** Cell cycle profile analysis of parental cells and rigosertib-resistant colonies treated with DMSO (light grey) or 100nM rigosertib (RGS, dark grey). Numbers within the plots indicate the percentage of cells with 2N or 4N DNA content. Representative graph of three experimental replicates. **(D)** Mitotic index quantification by phopho-Ser10 Histone H3 (p-HH3) staining in MDA-MB-453 parental cells (P, pink bars) and rigosertib resistant colonies (R, green bars) under 100nM and 500nM rigosertib, or DMSO as a control. Two-way ANOVA (mixed model) with Tukey multiple comparisons test: *p*<0.001 (***) and *p*<0.0001 (****). Representative graph of two experimental replicates. **(E and F)** Waterfall plot showing the enrichment (log2 fold change) of sgRNAs coding for WNK1 (**E**), and total gene enrichment (**F**), obtained from deep sequencing of the pooled resistant cells. **(G)** Volcano plot showing the differential gene expression (DGE) obtained by RNA-seq, comparing four resistant colonies versus the parental MDA-MB-453 cells. Blue dots represent significantly downregulated genes, whereas red dots show upregulated genes. **(H)** Immunoblot showing the WNK1 protein reduced expression levels in rigosertib-resistant colonies (R), compared to the MDA-MB-453 parental cells.

We then tested the differential enrichment of the sgRNA library in the pooled resistant cells, comparing it to the library with Cas9 activated by doxycycline (DOX) and treated with DMSO, the parental library without Cas9 induction, and the DNA library preparation. The rigosertib-treated cells showed a strong decrease in total sgRNA reads, probably due to massive cell death. Interestingly, there was a significant enrichment in a few reads over the parental controls (Supp. Figure 1D). The sequence identification showed enrichment of sgRNAs coding for the gene WNK1 (With-No-Lysin [K] 1 kinase), with four WNK1 coding sgRNA, out of the five enclosed in the Yusa library, at the top of the waterfall plot (Figure 1E). Moreover, when evaluating the sgRNA enrichment by genes, the WNK1 kinase gene was the top hit (Figure 1F). The presence of WNK1 sgRNA was confirmed in the isolated resistant colonies, mainly detecting the insertion of the sgRNA_5, targeting exon 4, and the sgRNA_2 which targets exon 2 of the WNK1 gene. WNK1 CRISPR edition was verified by T7 endonuclease activity assays (Supp. Figure 1E), and by sequencing the WNK1 locus indels generated (Supp. Figure 1F). Worth mentioning, we also found the WNK1 wild-type allele, suggesting that cells cannot completely eliminate WNK1 expression, as it is considered a common essential gene (Supp. Figure 1G). Indeed, the resistant colonies showed variability in their proliferative index compared to the parental cells (Supp. Figure 1H), yet they grew efficiently and the proliferative signaling downstream of WNK1 is not impaired, as seen by ERK5 MAP kinase activation (Supp. Figure 1I). Finally, we confirm the downregulation of WNK1 expression by RNA-seq in three of the isolated colonies compared to the parental cell transcriptome (Figure 1G), which was also reflected by a strong reduction in WNK1 protein levels (Figure 1H).

### 3.2. WNK1 depletion leads to rigosertib treatment resistance

To verify that WNK1 downregulation was responsible for the rigosertib-acquired resistance, we performed specific WNK1 depletion, by CRISPR-Cas9 edition (Supp. Figure 2A, B) in the MDA-MB-453 and MCF-7 cell lines, as seen by mRNA qRT-PCR (Figure 2A) and protein expression (Figure 2B). WNK1 knocked-down cells (WNK1-KD) have a slightly delayed proliferation rate compared to WNK1-WT cells (Supp. Figure 2C), yet not impairing cell growth. We then challenged the WNK1-KD cells to grow in the presence of rigosertib, observing increased survival when compared to WNK1-WT cells (Figure 2C, D). We obtained similar data by WNK1 shRNA downregulation (Supp. Figure 2D), as well in other cell lines such as EVSA-T and MDA-MB-231 by CRISPR-Cas9 depletion (Supp. Figure 2E). All these results were corroborated by cell competition assays, mixing WNK1-KD cells (EGFP labeled) and WNK1-WT cells (mRuby3 labeled) in a 20/80% ratio and culturing them in the presence of rigosertib (Supp. Figure 2G). DMSO-treated WNK1-KD cells showed a slight reduction in proliferation compared to WNK1-WT cells, whereas under rigosertib treatment, WNK1-KD cells outgrew the culture in approximately one week (MDA-MB-453) to 10 days (MCF-7) and were the only cells detected at the end of the experiment, as almost all WNK1-WT cells were dead (Figure 2E, F). Furthermore, chemical inhibition of WNK1 kinase activity, using WNK-463 and WNK-IN-11 inhibitors, also protected cells against rigosertib treatment (Figure 2G), indicating that WNK1 kinase activity is important for efficient cell response to rigosertib.

**Figure 2.**
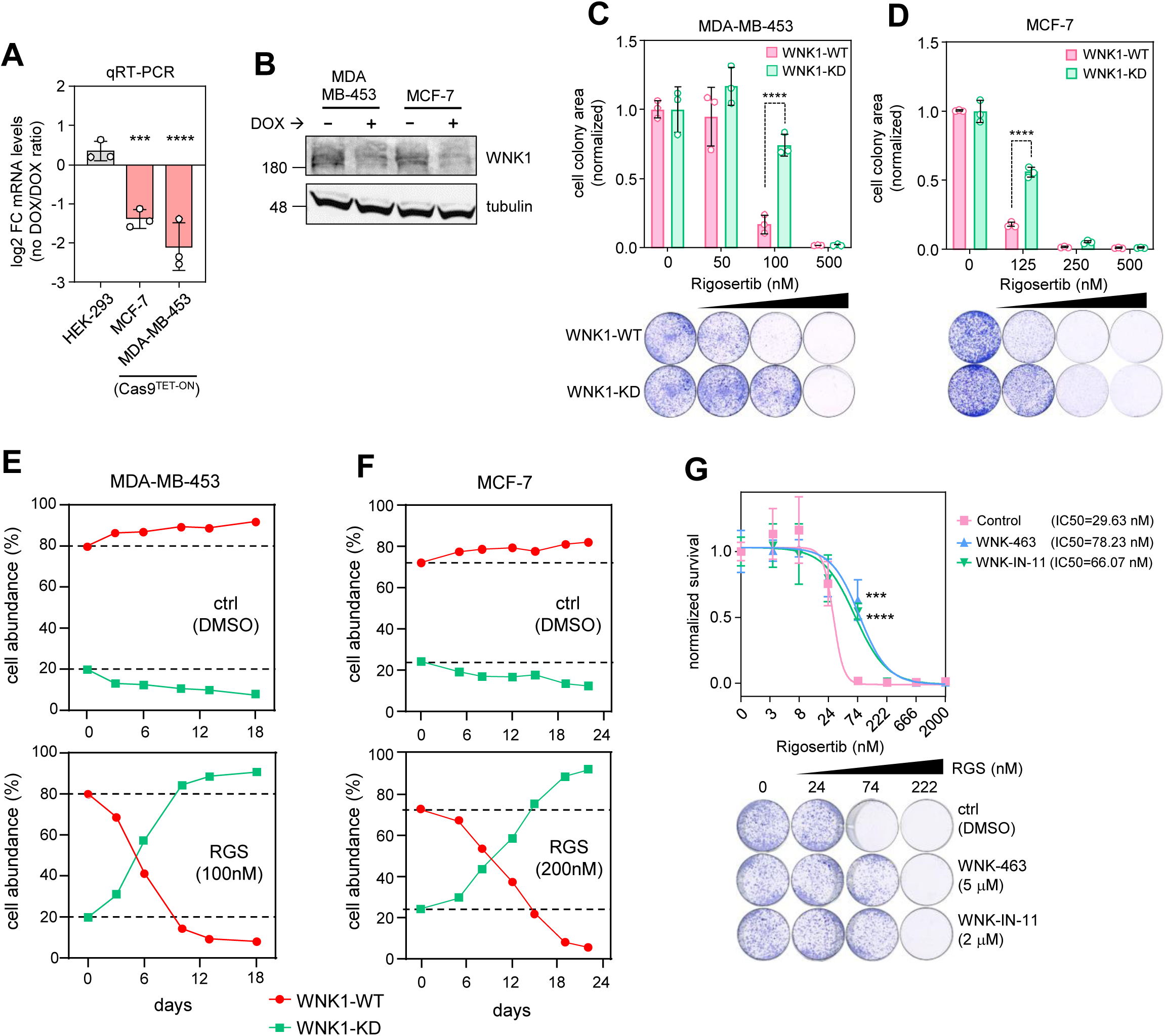
WNK1 inactivation leads to rigosertib resistance: **(A)** WNK1 mRNA expression by qRT-PCR in MCF-7 and MDA-MB-453 cells upon doxycycline (DOX) inducible Cas9 activation (Cas9^TET-ON^). HEK-293 were used as control cells. The plot represents the log2 fold chance ratio of no DOX versus DOX samples. One-way ANOVA with Dunnett’s multiple comparison test: *p*<0.001 (***) and *p*<0.0001 (****). Representative graph of three experimental replicates. **(B)** Immunoblot showing the reduction of WNK1 protein expression upon genetic depletion (+DOX) in MDA-MB-453 and MCF-7 cell lines. **(C and D)** Colony formation assay of MDA-MB-453 (**C**) and MCF-7 (**D**) WNK1-WT cells (pink bars) and WNK-KD cells (green bars) in the presence of different concentrations of rigosertib. Two-way ANOVA with Tukey’s multiple comparisons test: *p*<0.0001 (****). Representative graph of three experimental replicates. **(E and F)** Competition assays of MDA-MB-453 (**E**) and MCF-7 (**F**) WNK1-WT cells (red line) and WNK1-KD cells (green line) under DMSO (top panels) or rigosertib treatment (bottom panels) at the indicated concentrations. **(G)** Colony formation assay and normalized logarithmic regression IC50 calculation of MDA-MB-453 parental cells in the presence of the WNK1 inhibitors WNK-463 (blue line) and WNK-IN-11 (green line), or DMSO as control (pink line), and confronted to serial dilutions of rigosertib. Two-way ANOVA test: *p*<0.001 (***) and *p*<0.0001 (****). Representative graph of two experimental replicates

Finally, we confirmed that this effect of WNK1 downregulation is replicated in an in vivo context by implanting the MDA-MB-453 WNK1-WT and WNK1-KD cells into nude mice and treating them with either rigosertib sodium salt or vehicle (Figure 3A). This rigosertib dosing regimen was not toxic, as measured by animal weight and blood cell population counting (Supp. Figure 3A, B), yet it stopped WNK1-WT tumor growth soon after administration, whereas WNK1-KD tumors were still able to grow (Figure 3B). Histological analysis of tumors showed a reduction in the mitotic index in WNK1-KD tumors treated with rigosertib (Figure 3C), confirming the in vitro data from the resistant colonies (Figure 1D), and a reduction in cleaved caspase-3 staining, when compared to WNK1-WT tumors (Figure 3D), indicating cell survival.

**Figure 3.**
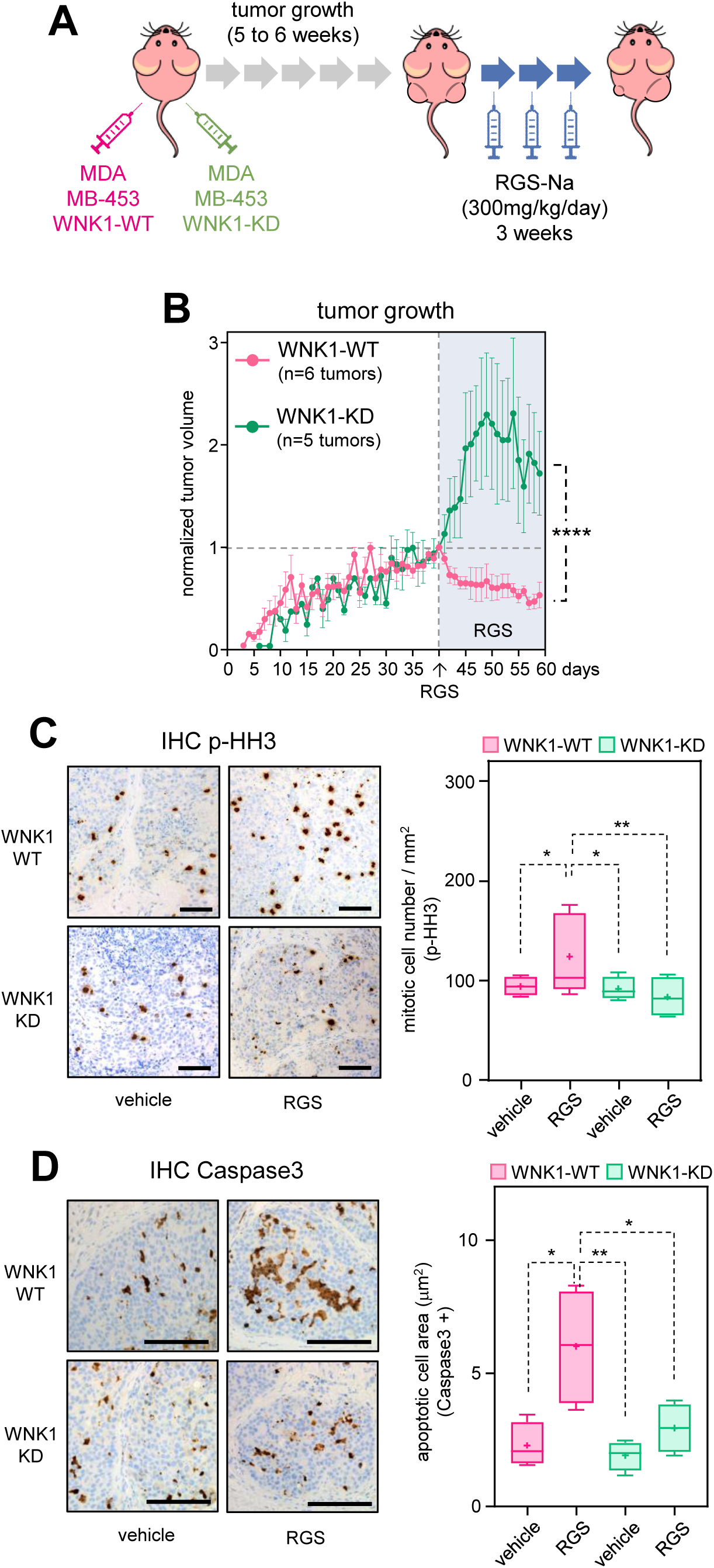
WNK1 inactivation provides survival against rigosertib during in vivo tumoral growth: **(A)** Cartoon of the xenograft experimental design. MDA-MB-453 WNK1-WT and WNK-KD are injected in the left and right flank of nude mice respectively. Once tumors reached approximately 100 mm^3^ (5 to 6 weeks), mice were treated with vehicle or rigosertib sodium salt (RGS-Na). **(B)** Tumor volume growth graph of both WNK1-WT (pink line) and WNK1-KD (green line). Tumor volume is normalized on the day of rigosertib administration (blue-shaded area). Two-way ANOVA comparison test of tumoral growth upon rigosertib treatment: *p*<0.0001 (****) **(C)** Mitotic index detection by immunohistochemistry staining of phospho-Ser10 Histone H3 (p-HH3) (left panel) (bar=100 μM), and percentage of p-HH3 positive cells quantification (right panel) in WNK1-WT (pink bars) and WNK1-KD (green bars) tumors upon rigosertib (RGS) treatment. Two-way ANOVA with Tukey’s multiple comparison test of four different tumoral samples per condition: *p*<0.05 (*), *p*<0.01 (**) **(D)** Cell death detection by immunohistochemistry staining of cleavage caspase 3 (left panel) (bar=100 μM), and quantification of caspase 3 positive signal area (right panel) in WNK1-WT (pink bars) and WNK1-KD (green bars) tumors upon rigosertib (RGS) treatment. Two-way ANOVA with Tukey’s multiple comparison test of four different tumoral samples per condition: *p*<0.05 (*), *p*<0.01 (**)

All of these data demonstrate that WNK1 kinase inactivation mediates acquired resistance to the mitotic drug rigosertib in vitro and in vivo.

### 3.3. WNK1 inactivation impedes mitotic arrest upon rigosertib treatment

Rigosertib is considered a mitotic drug (36–38), and WNK1 kinase has also been shown to modulate mitotic progression, although the underlying mechanism is still unknown (39). Thus, we studied the impact of rigosertib and WNK1 inactivation during mitotic progression. WNK1 genetic depletion led to a reduction in mitotic arrest upon rigosertib treatment, both in MDA-MB-453 and MCF-7 cells (Figure 4A, B) and in MDA-MB-231 cells (Supp. Figure 2F). In addition, WNK1 chemical inhibition rendered similar results (Figure 4C, D), indicating that WNK1 kinase activity impedes the mitotic arrest derived from rigosertib action.

**Figure 4.**
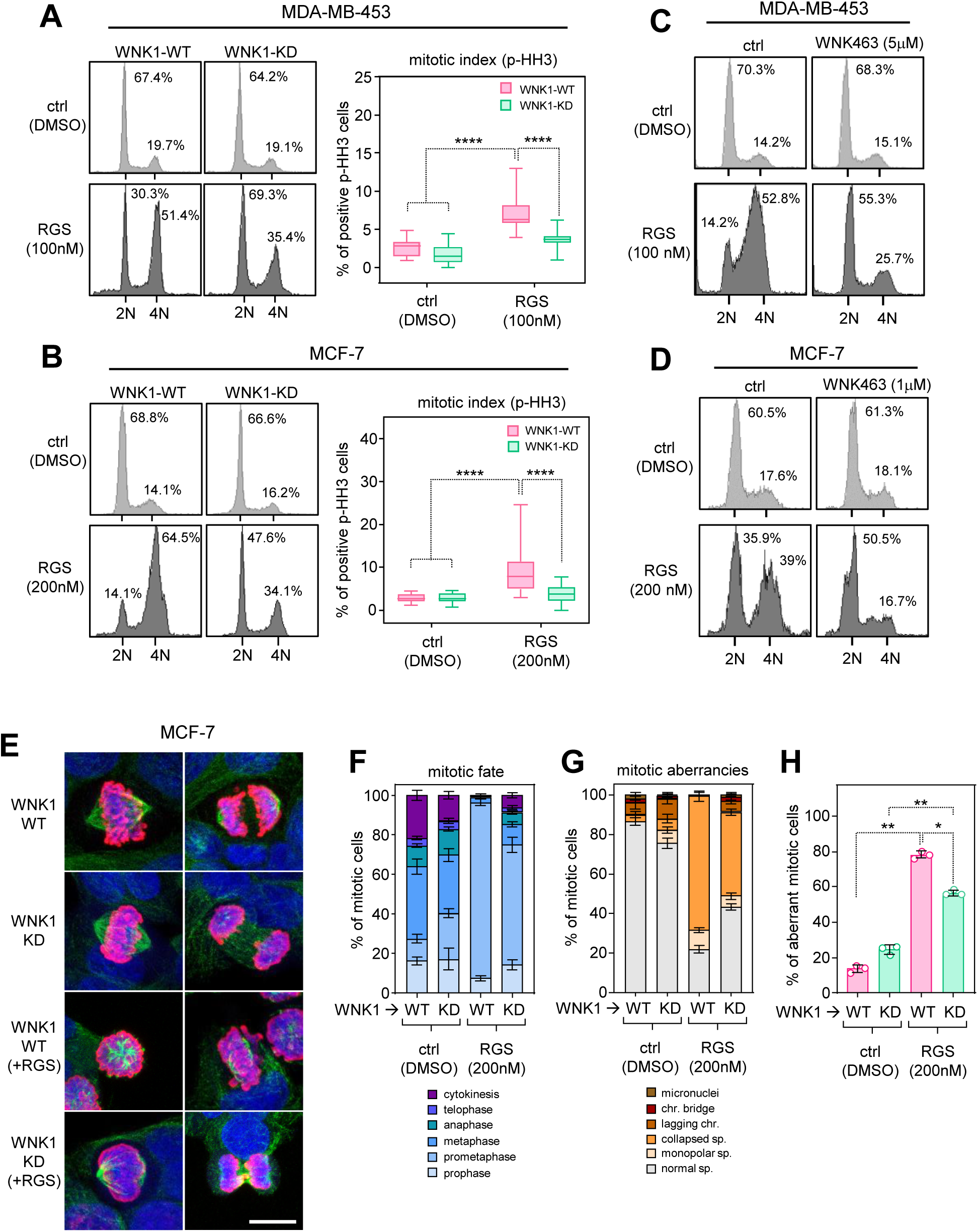
WNK1 inactivation impedes the mitotic arrest mediated by rigosertib. **(A and B)** Cell cycle profile analysis and mitotic index quantification by phospho-Ser10 Histone H3 (p-HH3) of MDA-MB-453 (**A**) and MCF-7 (**B**) WNK1-WT cells (pink bars) and WNK1-KD cells (green bars) treated with rigosertib or DMSO as control. Two-way ANOVA comparison test: *p*<0.0001 (****). Representative panel of three experimental replicates **(C and D)** Cell cycle profile analysis of MDA-MB-453 (**A**) and MCF-7 (**B**) WNK1-WT cells incubated with the WNK1 inhibitor WNK-463 and treated rigosertib or DMSO as control. Representative panel of three experimental replicates. **(E to H)** Mitotic fate study by confocal microscopy of WNK1-WT and WNK1-KD MCF-7 cells, in the presence of rigosertib, stained for alpha-tubulin (green), phospho-Ser10 histone H3 (red) and DAPI for DNA (blue) (**E**). Over 100 mitotic cells in each condition were evaluated for quantification of the different mitotic phases (**F**), and mitotic aberrations (**G**). Quantification of the total mitotic aberrancies for two-way ANOVA with Tukey’s multiple comparison test analysis: *p*<0.05 (*), *p*<0.01 (**) (**H**)

A detailed analysis of mitotic progression in MCF-7 WNK1-KD cells (Figure 4E) showed that there is a slight but not significant increase in mitotic aberrations compared to WNK1-WT cells (Figure 4H), not enough to induce mitotic arrest, and that all phases of mitotic progression can be detected (Figure 4F). On the other hand, when WNK1-WT cells are treated with rigosertib, there is a dramatic change in the fate of mitotic cells, with almost all cells stacked at the prometaphase stage, with collapsed spindles that are unable to properly align chromosomes, and with no visible advanced phases of mitosis such as anaphase or telophase (Figure 4F, G). These mitotic aberrations were alleviated in WNK1-KD cells, where we found cells in anaphase and telophase, suggesting that WNK1 inactivation reduces the toxic effect of rigosertib on the mitotic spindle.

### 3.4. Inactivation of WNK1 leads to microtubule stabilization, modifying the response to microtubule-associated drugs

Rigosertib is proposed to be a microtubule-related drug, binding to the α/β-tubulin dimer in the colchicine site, at the interphase of both tubulin monomers. This leads to a microtubule depolymerization causing mitotic progression impairment (40, 41). We, therefore, evaluated if WNK1 depletion could also modified the response to other microtubule depolymerizing drugs such as ABT-751, colchicine, or nocodazole. WNK1-KD cells had a significantly reduced cell cycle arrest in the 4N DNA content, upon the microtubule depolymerizing drugs treatment, when compared to WNK1-WT cells (Figure 5A), and this was also reflected in the capacity of WNK1-KD cells to outcompete the WNK1-WT cells when cell proliferation was evaluated by competition assays (Figure 5B). This indicates that WNK1 inactivation provides survival capacity to other microtubule poisons and it is not a specific mechanism for rigosertib. We also observed a similar response in the rigosertib-resistant colonies isolated from the screening process (Supp. Figure 4A, C, D). Of note, the impairment response to microtubule depolymerizing drugs was not due to the Leu240Phe mutation in the beta-tubulin gene (TUBB) (40) (Supp, Figure 4E).

**Figure 5.**
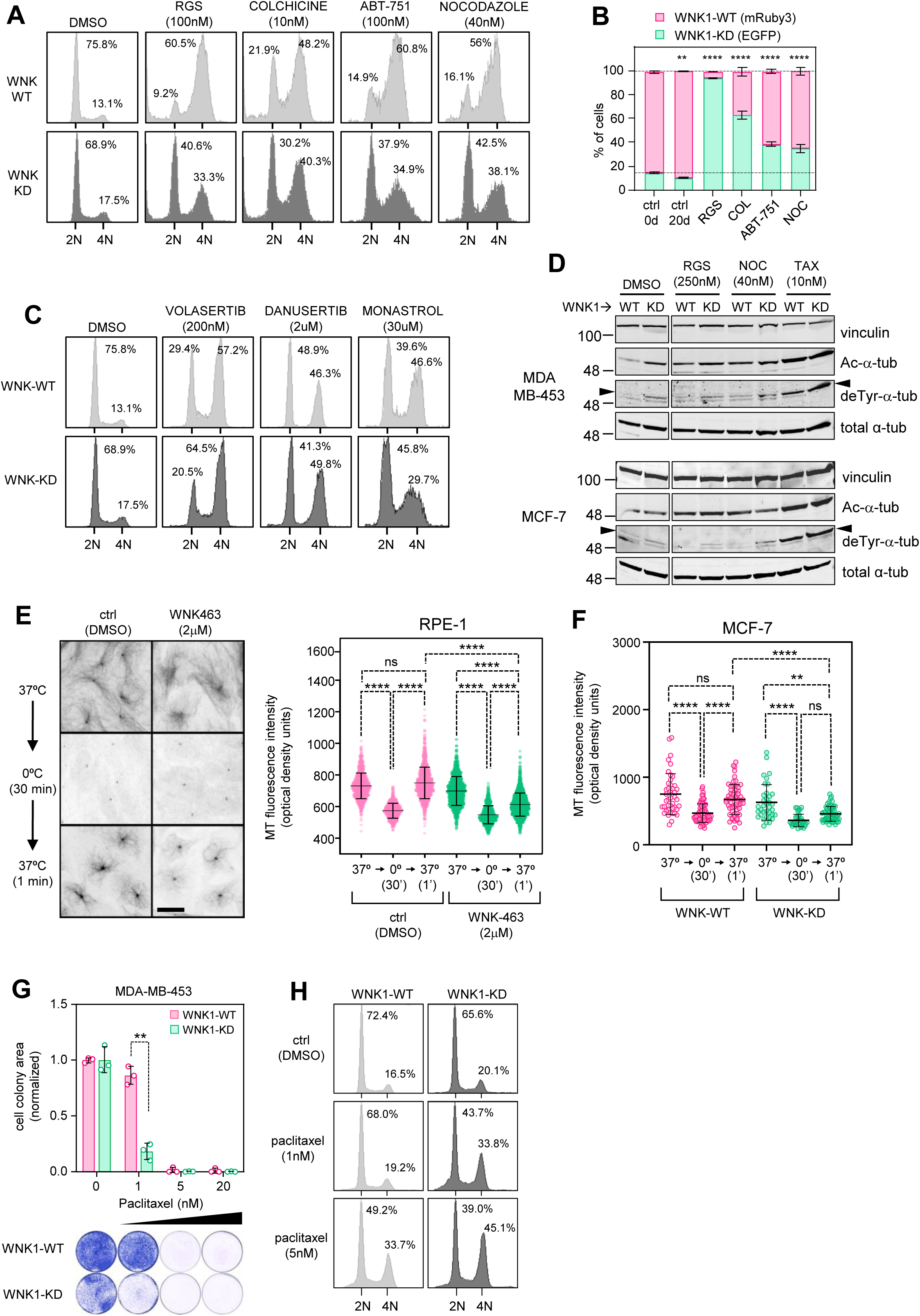
WNK1 inactivation influences the response to microtubule-related drugs: **(A)** Cell cycle profile of MDA-MB-453 WNK1-WT (light grey) and WNK1-KD cells (dark grey) treated with rigosertib (RGS), ABT-751, colchicine and nocodazole for 24h. Numbers within the plots indicate the percentage of cells in the 2N and 4N peaks. Representative panel of three experimental replicates. **(B)** Cell competition assay of MDA-MB-453 WNK1-WT (red bars) and WNK1-KD cells (green bars) incubated for 20 days in the presence of 100 nM rigosertib (RGS), 20 nM colchicine (COL), 100 nM ABT-751 and 25 nM nocodazole (NOC). Two-way ANOVA comparison test of the EGFP cell population: *p*<0.01 (**) and *p*<0.0001 (****). Representative graph of two experimental replicates. **(C)** Cell cycle profile of MDA-MB-453 WNK1-WT (light grey) and WNK1-KD cells (dark grey) treated with rigosertib volasertib, danusertib and monastrol for 24 hours. Numbers within the plots indicate the percentage of cells in the 2N and 4N peaks. Representative graph of three experimental replicates **(D)** Biochemical analysis of tubulin stabilization markers by immunoblotting for acetylated alpha-tubulin (Ac-α-tub) and detyrosinated alpha-tubulin (deTyr-α-tub) in MDA-MB-453 (top panel) and MCF-7 (bottom panel) WNK1-WT and WNK1-KD cells treated with rigosertib (RGS), nocodazole (NOC) and paclitaxel (TAX) for 6 hours. Total alpha-tubulin and vinculin detection are used as loading controls. **(E)** Alpha-tubulin staining images of the microtubule polymerization assay by 30 min of cold shock and 1 min recovery in warm media (left panel), and alpha-tubulin intensity quantification plot in in RPE-1 cells treated with WNK-463 (green dots) or DMSO as control (pink dots) (right panel). Bar = 20 μM. One-way ANOVA with Tukey’s comparison test: *p*<0.0001 (****), and not significant (ns). **(F)** Microtubule polymerization assay quantification in MCF-7 WNK1-WT (pink dots) and WNK-KD cells (green dots), done over 100 cells in each condition. One-way ANOVA with Tukey’s comparison test: *p*<0.01 (**), *p*<0.0001 (****), and not significant (ns). **(G)** Colony formation assays of WNK1-WT (red bars) and WNK1-KD (green bars) cells incubated in different concentrations of paclitaxel. Two-way ANOVA with Tukey’s multiple comparison test: *p*<0.01 (**). Representative graph of two experimental replicates. **(H)** Cell cycle profile of MDA-MB-453 WNK1-WT (light grey) and WNK1-KD cells (dark grey) treated with paclitaxel for 24 h. Numbers within the plots indicate the percentage of cells in the 2N and 4N peaks.

Since microtubule-related drugs cause severe mitotic arrest, we wondered whether WNK1 inactivation might synergize with any drug that causes mitotic arrest, regardless of its mechanism of action. Therefore, we tested the response of WNK1-KD cells to other mitotic inhibitors (Figure 5C). Volasertib (Plk1 kinase inhibitor) and danusertib (Aurora A/B kinase inhibitor) induced a similar 4N cell cycle arrest in WNK1-WT and WNK1-KD cells, indicating that a mitotic arrest context is not the cause of the differential response of WNK1-KD cells to microtubule poisons. Surprisingly, the Eg5 kinesin inhibitor monastrol resulted in reduced cell cycle arrest in WNK1-KD cells. Eg5 kinesin is not only essential for centrosome separation and mitotic spindle assembly (42), but is also involved in controlling microtubule dynamics (43–45), explaining the reduction of cell cycle arrest in WNK1-KD cells, and suggesting that changes in microtubule dynamics are key for the protective effect of WNK1 inactivation against microtubule poisons.

To investigate if this was the case, we tested whether WNK1 inactivation could alter the stability of microtubule filaments. MDA-MB-453 and MCF-7 WNK1-KD cells showed elevated levels of acetylated (Ac-Tub) and detyrosinated (deTyr-Tub) tubulin, two post-traductional modifications concordant with increased stability of the microtubule filaments (46–48) (Figure 5D). Noteworthy, the augmented levels of Ac-Tub and deTyr-Tub were also present in WNK1-KD cells treated with rigosertib or nocodazole, while paclitaxel addition led to a strong signal of both tubulin modifications due to massive stabilization of the microtubules. This suggests that WNK1 inactivation can modify the microtubule dynamics by making them more stable, thus lessening the response to microtubule depolymerizing drugs such as rigosertib and others. To verify this hypothesis, we performed functional microtubule dynamic assays, by testing the polymerization capacity of tubulin (49). RPE-1 cells were incubated with WNK1 inhibitor WNK-463, or DMSO as control, then the microtubule network was depolymerized by cold treatment and allowed to repolymerize in warm media (Figure 5E). WNK1-inhibited RPE-1 cells showed less microtubule nucleation capacity. The same data was obtained in the MCF-7 cells genetically depleted for WNK1 (Figure 5F), confirming that WNK1 inactivation alters microtubule polymerization dynamics.

As WNK1 inactivation modulates microtubule dynamics by increasing their stability, we wondered if this could also provide a differential response to microtubule-stabilizing drugs such as paclitaxel. Firstly, we tested the survival capacity of the isolated rigosertib-resistant colonies to paclitaxel, observing a significant reduction in survival compared to parental cells (Supp, Figure 4D). This differential response to paclitaxel was dependent on WNK1 depletion since WNK1-KD cells were more sensitive to paclitaxel than WNK1-WT cells (Figure 5G). This data is concurrent to the increased 4N cell cycle arrest in WNK1-KD cells (Figure 5H), demonstrating that WNK1 inactivated cells have more stable microtubules and therefore need less amount of paclitaxel to achieve a cytotoxic effect.

Overall, our data suggest that the WNK1 kinase modulates microtubule dynamics and that WNK1 inactivation confers resistance to microtubule depolymerizing drugs (rigosertib, colchicine, nocodazole) and susceptibility to microtubule stabilizing drugs such as taxanes.

### 3.5. Osmotic stress leads to a differential response of microtubule-associated drugs

WNK1 is a major controller of cellular ion homeostasis, being essential for cells to recover from an osmotic stress (14). In addition, osmotic stress has been shown to alter the cell cycle and mitotic progression (25, 50). Therefore, we interrogated if osmotic stress provides a differential response to rigosertib while cells are grown in different osmotic conditions (Figure 6A, B). Hypoosmotic stress enhanced the rigosertib-mediated 4N cell cycle arrest in WNK1-WT cells when compared to isoosmotic conditions. Importantly, WNK1-KD cells had a significantly reduced cell cycle arrest (Figure 6A), and better survival (Figure 6B) in both isoosmotic and hypoosmotic conditions, indicating that WNK1 controls the response to rigosertib by counteracting the osmotic stress signaling. Interestingly, cells under hyperosmotic stress did not show any cell cycle arrest upon rigosertib (Figure 6A), correlating with increased cell survival (Figure 6B), thus mimicking the inactivation of WNK1.

**Figure 6.**
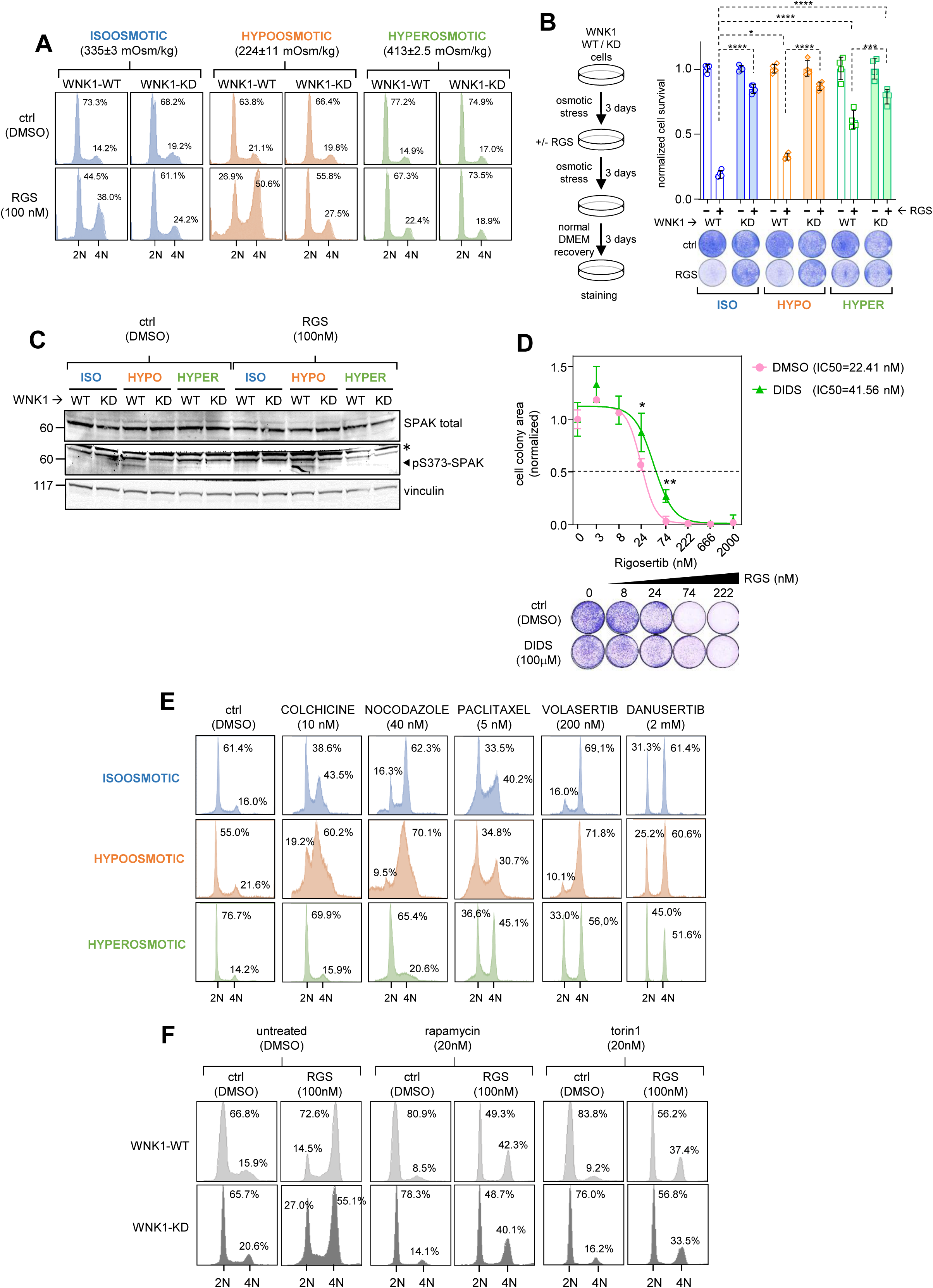
Osmotic stress leads to differential response to microtubule-related drugs: **(A)** Cell cycle profile of MDA-MB-453 WNK1-WT and WNK1-KD cells treated with rigosertib (RGS) under isoosmotic (blue profiles), hypoosmotic (orange profiles) and hyperosmotic conditions (green profiles). The average of three osmolarity measurements (±SD) are indicated in mOsm/kg. Numbers within the plots show the percentage of cells in the 2N and 4N peaks. Representative panel of three experimental replicates. **(B)** Colony formation assay of MDA-MB-453 WNK1-WT and WNK1-KD cells treated with rigosertib (RGS) under different osmotic media conditions. Histogram shows the quantification of cell survival under isoosmotic (blue bars), hypoosmotic (orange bars) and hyperosmotic conditions (green bars). Two-way ANOVA with Tukey’s multiple comparison test: *p*<0.05 (*), *p*<0.001 (***), *p*<0.0001 (****). Representative graph of two experimental replicates. **(C)** WNK1 signaling analysis by western blot of MDA-MB-453 WNK1-WT and WNK1-KD cells incubated under isoosmotic, hypoosmotic, and hyperosmotic conditions for three days, combined with rigosertib (RGS) or DMSO as control. Antibodies for total SPAK and phospho-SPAK-Ser373 residue (arrowhead) were probed. (*) labels non-specific cross reactivity band. Vinculin expression is used as loading control **(D)** Colony formation assay and IC50 calculation of MDA-MB-453 cells grown in the presence of the chloride transport inhibitor DIDS (green line), or DMSO as control (pink line), and tested against serial dilutions of rigosertib (RGS). Two-way ANOVA with Tukey’s multiple comparison test: *p*<0.05 (*) and *p*<0.01 (**). Representative graph of two experimental replicates. **(E)** Cell cycle profile analysis of MDA-MB-453 cells incubated under isoosmotic (blue profiles), hypoosmotic (orange profiles) and hyperosmotic (green profiles), treated with the indicated drugs. Numbers within the plots show the percentage of cells in the 2N and 4N peaks. **(F)** Cell cycle profile analysis of MDA-MB-453 WNK1-WT (light grey) and WNK1-KD (dark grey) cells incubated in with the mTORC inhibitors rapamycin and torin1, and then tested for cell cycle arrest with rigosertib (RGS). Numbers within the plots show the percentage of cells in the 2N and 4N peaks.

WNK1 is known to be activated under hypertonic stress (51, 52), and also under hypotonic stress (53, 54). All these data are mostly retrieved in short-term osmotic stress experiments (minutes), whereas our osmotic stress assays were done for longer times (days). Thus, we evaluated WNK1 activation signaling under longer osmotic stress induction, by testing the phosphorylation of the WNK1 effector kinase SPAK (Figure 6C). In our hands, hypoosmotic stress led to the activating phosphorylation of SPAK in a WNK1-dependent manner, whereas neither hyperosmotic stress nor the presence of rigosertib altered SPAK phosphorylation. These data indicate that hypoosmotic stress activates WNK1 leading to increased sensitivity to rigosertib, while hyperosmotic stress protects against rigosertib by mimicking WNK1 inactivation. In addition, altering the flux of chloride ions, using the ClC-Ka chloride channel blocker DIDS (4,4’-diisothiocyanato-2,2’-stilbenedisulfonic acid disodium salt), also provided an increased survival response to rigosertib, demonstrating that chloride homeostasis is critical for the response to the microtubule destabilizing drug rigosertib (Figure 6D).

Osmotic stress modulates microtubule dynamics by modifying the cytoplasm viscosity and increasing the intracellular molecular crowding (55). Thus, we tested whether osmotic stress leads to differential response to other microtubule-related drugs (Figure 6E). Hypoosmotic stress provided increased sensitivity to microtubule depolymerizing compounds (colchicine and nocodazole) and a less efficient arrest to the stabilizing drug paclitaxel. On the other hand, hyperosmotic stress provided a complete protection to colchicine and nocodazole, whereas paclitaxel was more efficient in eliciting a 4N cell cycle arrest. Noteworthy, these changes in drug response were only related to microtubule drugs, since the 4N cell cycle arrest upon Plk1 inhibition (volasertib) or Aurora kinase inhibition (danusertib) did not lead to any differential response associated with osmotic stress. Only in the case of hyperosmotic stress, there was a reduction in the 4N arrest upon Plk1 or aurora kinases inhibition, but this can be explained by the delay in cell cycle progression under hypertonic stress (25), provoking fewer cells entering into mitosis.

Intracellular molecular crowding is not only controlled by WNK1 (52), but also modulated by mTOR signaling by adjusting ribosome concentration (56). For that reason, we tested if mTOR inhibition, and the subsequent molecular crowding reduction, could rescue the acquired resistance to rigosertib upon WNK1 depletion (Figure 6F). Although mTOR inhibition results in a significant G1 arrest (57, 58) and therefore rigosertib-mediated 4N arrest was less effective, WNK1-KD cells reached similar 4N arrest levels as WNK1-WT cells, suggesting that WNK1 inactivation increases molecular crowding, mediating the differential response to rigosertib.

## 4. DISCUSSION

Upon a CRISPR-Cas9 resistance screen for identification of rigosertib-associated biomarkers, we found that inactivation of the osmostress kinase WNK1 mediates cellular resistance to rigosertib (Figure 1). Using genetic depletion and chemical inhibition approaches, we validated WNK1 as a biomarker for rigosertib resistance through in vitro cell proliferation and competition assays, and in vivo xenograft tumor assays (Figures 2 and 3). Noteworthy, Jost and colleagues (40) also found WNK1 as a positive hit in their CRISPR screen for rigosertib response, as depicted from the Open Repository of CRISPR Screens in BioGRID database platform (https://orcs.thebiogrid.org/Screen/1176) (59).

We show that WNK1 inactivation does not modify targeted therapies such as volasertib (Plk1 inhibitor) or trametinib (MEK inhibitor) (Supp. Figure 1) reinforcing the idea that rigosertib does not act as a RAS-MEK or PLK1 inhibitor in our hands. On the contrary, we show that WNK1 inactivation leads to increased microtubule stability, thus not only conferring resistance to rigosertib, but to other similar microtubule poisons such as colchicine, ABT-751 or nocodazole (Figure 5), and confirming recent data on rigosertib’s primary mechanism of action as a microtubule destabilizer (40, 41, 60). Concomitantly, WNK1 depletion confers sensitivity to paclitaxel, a microtubule-stabilizing drug, demonstrating that WNK1 activity can modulate microtubule dynamics (Figure 5). The effect of WNK1 on microtubule dynamics may also explain why WNK1 silencing leads to mitotic aberrations (39), although we do not observe such a strong mitotic aberrant phenotype upon WNK1 depletion (Figure 4), indicating that WNK1 regulation of the mitotic spindle may be cell type-dependent. Importantly, we did not find the Leu240Phe TUBB mutation that confers rigosertib resistance (40), hinting a different molecular mechanism for altering tubulin dynamics upon WNK1 inactivation.

WNK1 is a master regulator for the osmotic stress response (14) and a sensor of molecular crowding induced by osmotic stress (52). The rigosertib-resistant colonies show not only strong downregulation of WNK1, but also of other ion homeostasis regulators such as the calcium-activated chloride channel ANO1 (TMEM16A) (61), annexin II light chain (S100A10) (62) or the lncRNAs NEAT1 and MALAT1, both known to be involved in molecular condensation and nuclear speckle crowding (63) (Figure 1), suggesting that rigosertib-resistant cells have undergone important changes in osmotic stress response signaling. Interestingly, osmotic stress has also been proposed to regulate microtubule dynamics by altering the intracellular molecular crowding and the cytoplasm physicochemical properties (55, 64). Thus, we evaluated if alterations in cytoplasmic tonicity and crowding, due to osmotic stress, could modify the response to microtubule-associated drugs (Figure 6). Whereas hypoosmotic stress leads to a more efficient response to microtubule depolymerizing drugs and reduces the efficacy of taxanes as stabilizing compounds, hyperosmotic stress has the opposite effect, confirming that imbalances in ion homeostasis can alter the response to microtubule-related drugs. Importantly, WNK1 inactivation ameliorates these alterations under hypoosmotic stress, having no impact in hyperosmotic conditions. Moreover, we can eliminate the differential response to rigosertib by reducing the molecular crowding using mTOR inhibitors, suggesting that WNK1 depletion mimics a hyperosmotic scenario leading to increased intracellular molecular crowding, thus altering the microtubule dynamics properties.

Hypoosmotic stress reduces the intracellular chloride levels, while hyperosmosis leads to elevated chloride ions. WNK1 is a chloride sensor that is autoinhibited upon chloride binding, and activated when chloride concentrations are low (65). Although WNK1 is classically proposed to be activated by hyperosmotic stress (51, 52, 66), hypotonic stress can also elicit WNK1 kinase activity (53, 54). Is important to mention that all these previous studies related to WNK1 activation are performed using short-term osmotic stress induction, while our cell cycle and cell survival assays are conducted over longer times. In our experimental setting, only hypoosmotic media activates WNK1 (by phospho-SPAK detection) whereas hypertonic stress has little effect (Figure 6). These data are consistent with the differential responses to microtubule-associated drugs observed upon osmotic stress induction and WNK1 depletion, confirming that WNK1 modulates microtubule dynamics in an osmoregulatory manner.

Overall, our data demonstrate that WNK1 activity is a biomarker for microtubule-associated drug response, and this effect is dependent on osmotic stress and the subsequent variations in the intracellular molecular crowding that modify the microtubule polymerization dynamics.

WNK1 has also been described to modulate the dynamics of the actin cytoskeleton (67–69), demonstrating that WNK1 can control the cytoskeleton in a variety of actions and suggesting that we cannot rule out other possible mechanisms. For example, the WNK1 downstream osmoregulator SGK1 kinase is known to be a modulator for microtubule dynamics, promoting microtubule depolymerization by phosphorylation of Tau in neurite formation (70). Whether this is a conserved mechanism in cancer epithelial cells is yet unknown. Important to mention, WNK1 abrogation leads to a very strong resistance to rigosertib, that it is not as efficient with other microtubule depolymerizing drugs with similar molecular mode if action like colchicine or ABT-751 (40) (Figure 5). This may be explained by the other possible molecular mechanisms behind rigosertib, especially as an inducer of oxidative stress by activating the JNK-mediated stress cascade (71), and by the fact that WNK1 is also a stress sensor that can modulate JNK signaling (72). Although we do not detect WNK1 activation upon rigosertib treatment, this is a possibility worth to explore in the future. Similarly, another report shows that the ability of rigosertib to destabilize microtubules is strongly inhibited by increased levels of uric acid (73). Although the authors here claim that uric acid competes for binding to tubulin, thereby inhibiting the effect of rigosertib, we cannot exclude the possibility of osmotic stress induction due to increased uric acid levels.

In conclusion, our data show that WNK1 kinase, through its function as an osmotic sensor, can modulate the response to anticancer microtubule-associated drugs. This is clinically relevant and worthy of further investigation in a more clinical scenario, as osmotic regulation is often altered in cancer patients and may therefore influence the efficacy of response to certain chemotherapeutic strategies.

## 5. HIGHLIGHTS

- WNK1 inactivation resulted in an impaired response to the mitotic drug rigosertib.
- This effect is mediated by an alteration in microtubule dynamics, confirming the predominant mechanism of rigosertib as a microtubule poison.
- WNK1 inactivation increases microtubule stability, reducing the efficacy of microtubule depolymerizing drugs and sensitizing to microtubule stabilizing agents.
- WNK1 modulates this mechanism by controlling osmotic stress and intracellular molecular crowding.

## 6. SIGNIFICANCE STATEMENT

Targeting cell division is an effective strategy to inhibit cancer proliferation. Microtubule inhibitors are compounds, which target mitosis, are a widely used therapeutic approach across various cancer types. However, their effectiveness is often compromised by molecular mechanisms of acquired resistance. Here we describe how osmotic stress and inactivation of the ion homeostasis kinase WNK1 can influence the response to microtubule-related drugs, conferring resistance or sensitivity depending on the drug’s mechanism of action. These findings raise a note of caution for cancer patients undergoing chemotherapeutic treatment with microtubule poisons, as disruptions in ion homeostasis may alter the therapeutic efficacy.

## Supporting information

Monfort-Vengut et al_Main Figures

## ACKNOWLEDGEMENTS

We thank Oscar Fernandez-Capetillo (CNIO) and Marcos Malumbres (CNIO) for sharing reagents, and Luis del Peso (IIBM-UAM) for helping with the RNA-seq analysis.

This study has been financed by the following grants: From the Ministerio de Ciencia, Innovación, Agencia Estatal de Investigación MCIN/AEI/FEDER UE (doi:10.13039/501100011033): RTI2018-095496-B-I00 and PID2021-125705OB-I00 (GdC); PID2021-122222OB-I00 (RSP); Juan de la Cierva Postdoctoral program FJC2020-044620-I (NSG). From the Spanish Association Against Cancer (AECC) Scientific Foundation: LABAE16017DECA (GdC) and POSTD234371SANZ (NSG). From the Spanish National Research Council (CSIC) with grants 201820I114, 2021-AEP035, and 2022-20I018 (GdC). From the UCLM with grant 2022-GRIN-34150 (RSP). EMBO Scientific Exchange Grant (SEG-9883) to AMV, in RS laboratory.

## 7. AUTHOR CONTRIBUTIONS

AMV performed the CRISPR screening and all the experimental procedures, with the help of BO, SB, AC, and AM. NSG performed the RNA-seq bioinformatic analysis and in vitro experiments. MR and RS participated in and supervised the tumor xenograft experiments. JMRR performed the NGS sequencing and analysis of the CRISPR libraries. JML and RSP provided intellectual input. GdC conceived and supervised the study. All authors participated in the data analysis, and GdC wrote the paper with the help of AMV and NSG.

