## Supplementary material for "Osmotic Stress Influences Microtubule Drug Response Via WNK1 Kinase Signaling": Monfort-Vengut et al_Main Figures

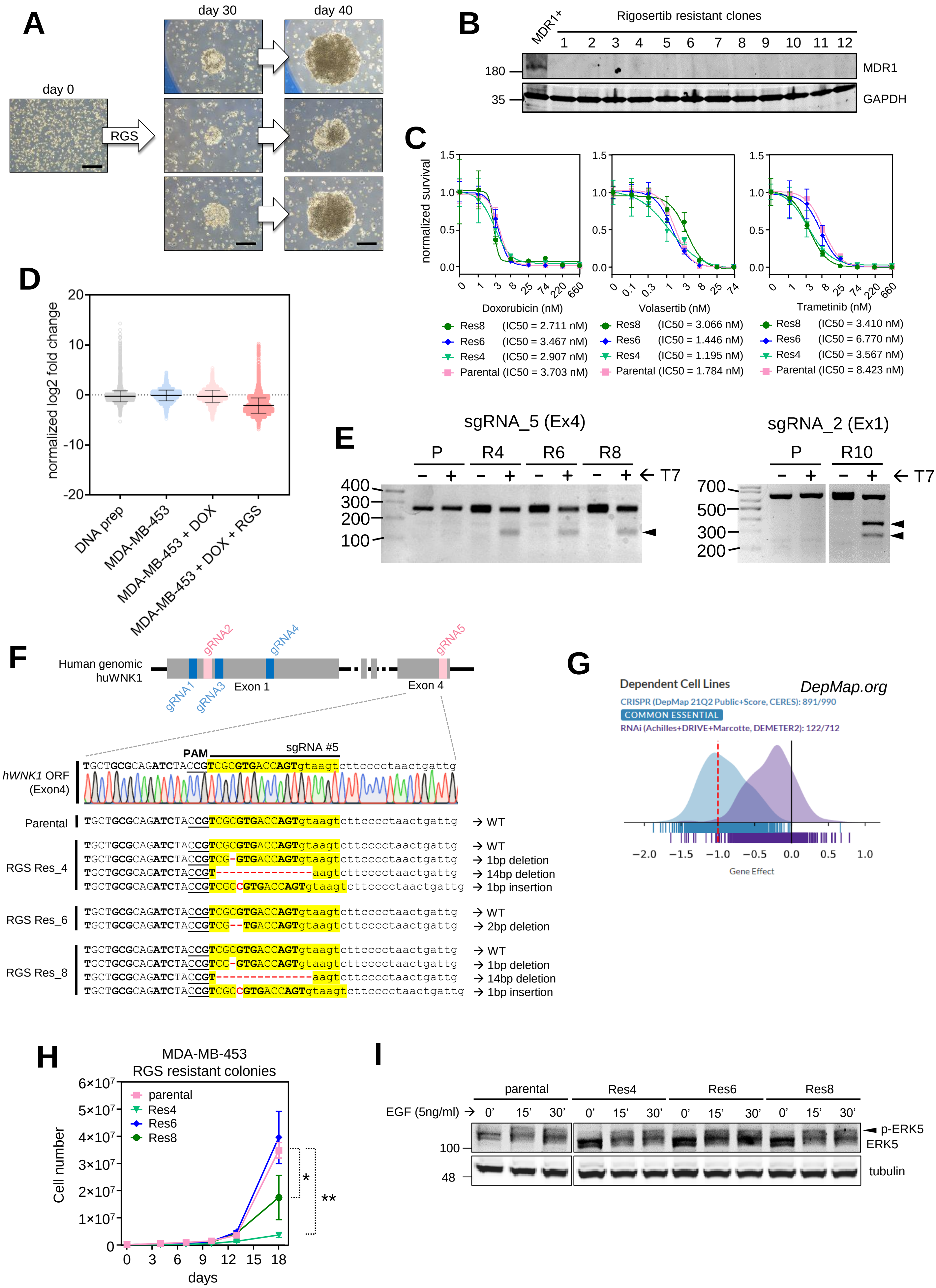
A
day 0
RGS
day 30
day 40
B
MDR1+
Rigosertib resistant clones
1 2 3 4 5 6 7 8 9 10 11 12
180
35
MDR1
GAPDH
C
normalized survival
Doxorubicin (nM)
Volasertib (nM)
Trametinib (nM)
Res8 (IC50 = 2.711 nM)
Res6 (IC50 = 3.467 nM)
Res4 (IC50 = 2.907 nM)
Parental (IC50 = 3.703 nM)
Res8 (IC50 = 3.066 nM)
Res6 (IC50 = 1.446 nM)
Res4 (IC50 = 1.195 nM)
Parental (IC50 = 1.784 nM)
Res8 (IC50 = 3.410 nM)
Res6 (IC50 = 6.770 nM)
Res4 (IC50 = 3.567 nM)
Parental (IC50 = 8.423 nM)
D
normalized log2 fold change
DNA prep
MDA-MB-453
MDA-MB-453 + DOX
MDA-MB-453 + DOX + RGS
E
sgRNA_5 (Ex4)
sgRNA_2 (Ex1)
P R4 R6 R8
P R10
T7
F
Human genomic huWNK1
gRNA1 gRNA2 gRNA3 gRNA4 gRNA5
Exon 1
Exon 4
PAM
sgRNA #5
hWNK1 ORF (Exon4)
TGCTGCGCAGATCTACCGTCGCGTGACCAGTgtaagtcttcccctaactgattg
Parental
TGCTGCGCAGATCTACCGTCGCGTGACCAGTgtaagtcttcccctaactgattg → WT
RGS Res_4
TGCTGCGCAGATCTACCGTCGCGTGACCAGTgtaagtcttcccctaactgattg → WT
TGCTGCGCAGATCTACCGTCG-GTGACCAGTgtaagtcttcccctaactgattg → 1bp deletion
TGCTGCGCAGATCTACCGT--------------aagtcttcccctaactgattg → 14bp deletion
TGCTGCGCAGATCTACCGTCGCCGTGACCAGTgtaagtcttcccctaactgattg → 1bp insertion
RGS Res_6
TGCTGCGCAGATCTACCGTCGCGTGACCAGTgtaagtcttcccctaactgattg → WT
TGCTGCGCAGATCTACCGTCG--TGACCAGTgtaagtcttcccctaactgattg → 2bp deletion
RGS Res_8
TGCTGCGCAGATCTACCGTCGCGTGACCAGTgtaagtcttcccctaactgattg → WT
TGCTGCGCAGATCTACCGTCG-GTGACCAGTgtaagtcttcccctaactgattg → 1bp deletion
TGCTGCGCAGATCTACCGT--------------aagtcttcccctaactgattg → 14bp deletion
TGCTGCGCAGATCTACCGTCGCCGTGACCAGTgtaagtcttcccctaactgattg → 1bp insertion
G
Dependent Cell Lines
DepMap.org
CRISPR (DepMap 21Q2 Public+Score, CERES): 891/990
COMMON ESSENTIAL
RNAi (Achilles+DRIVE+Marcotte, DEMETER2): 122/712
Gene Effect
H
MDA-MB-453
RGS resistant colonies
parental
Res4
Res6
Res8
Cell number
days
I
parental Res4 Res6 Res8
EGF (5ng/ml)
0' 15' 30'
p-ERK5
ERK5
tubulin
100
48

**SUPPLEMENTARY FIGURE LEGENDS**

Supplementary Figure 1

Rigosertib CRISPR-Cas9 screen and resistant colonies characterization:

(A) Representative pictures of the MDA-MB-453 cells before rigosertib addition (day 0), resistant colony detection (day 30) and growth (day 40). Bar = 200 $\mu$M

(B) MDR1 protein detection in the isolated MDA-MB-453 rigosertib resistant colonies. Positive control for MDR1 are MDA-MB-453 cells resistant to the Plk1 inhibitor volasertib.

(C) Colony formation and IC50 calculation, of MDA-MB-453 parental cells and rigosertib resistant cells, in the presence of doxorubicin, volasertib and trametinib.

(D) Analysis of the sgRNA number variations in the CRISPR library in different steps of the library preparation and the screening process. “DNA prep” is the plasmid format library. “MDA-MB-453” is the library already inserted and sorted in the MDA-MB-453 cell line. “MDA-MB-453 + DOX” is the sgRNA library upon Cas9 activation for 3 days. “MDA-MB-453 + DOX + RGS” is the sgRNA library after Cas9 activation and 40 days of rigosertib selection.

(E) T7-endoculease assays of parental MDA-MB-453 (P) and three resistant colonies (R4, R6 and R8) demonstrating the presence of WNK1 edition by the sgRNA_5 targeting WNK1 exon 4 (left panel), and R10 harboring the sgRNA_2 that targets WNK1 exon 1 (right panel).

(F) DNA sequencing of the WNK1 locus at the exon 4, in parental and rigosertib (RGS) resistant cells (Res4, Res6 and Res8), showing the indels generated by the WNK1 sgRNA_5.

(G) Cell dependency plot of WNK1 CRISPR depletion (blue peak) or WNK1 shRNA silencing (purple peak) obtained from the DepMap platform (www.depmap.org).

(H) Cell growth curve of parental and rigosertib resistant MDA-MB-453 cells. Two-way ANOVA test: $p$<0.05 (*) and $p$<0.01 (**)

(I) ERK5 activation detection in parental and rigosertib resistant MDA-MB-453 cells by immunoblotting for phospho-ERK5 (arrowhead) upon EGF induction at the indicated times (minutes).

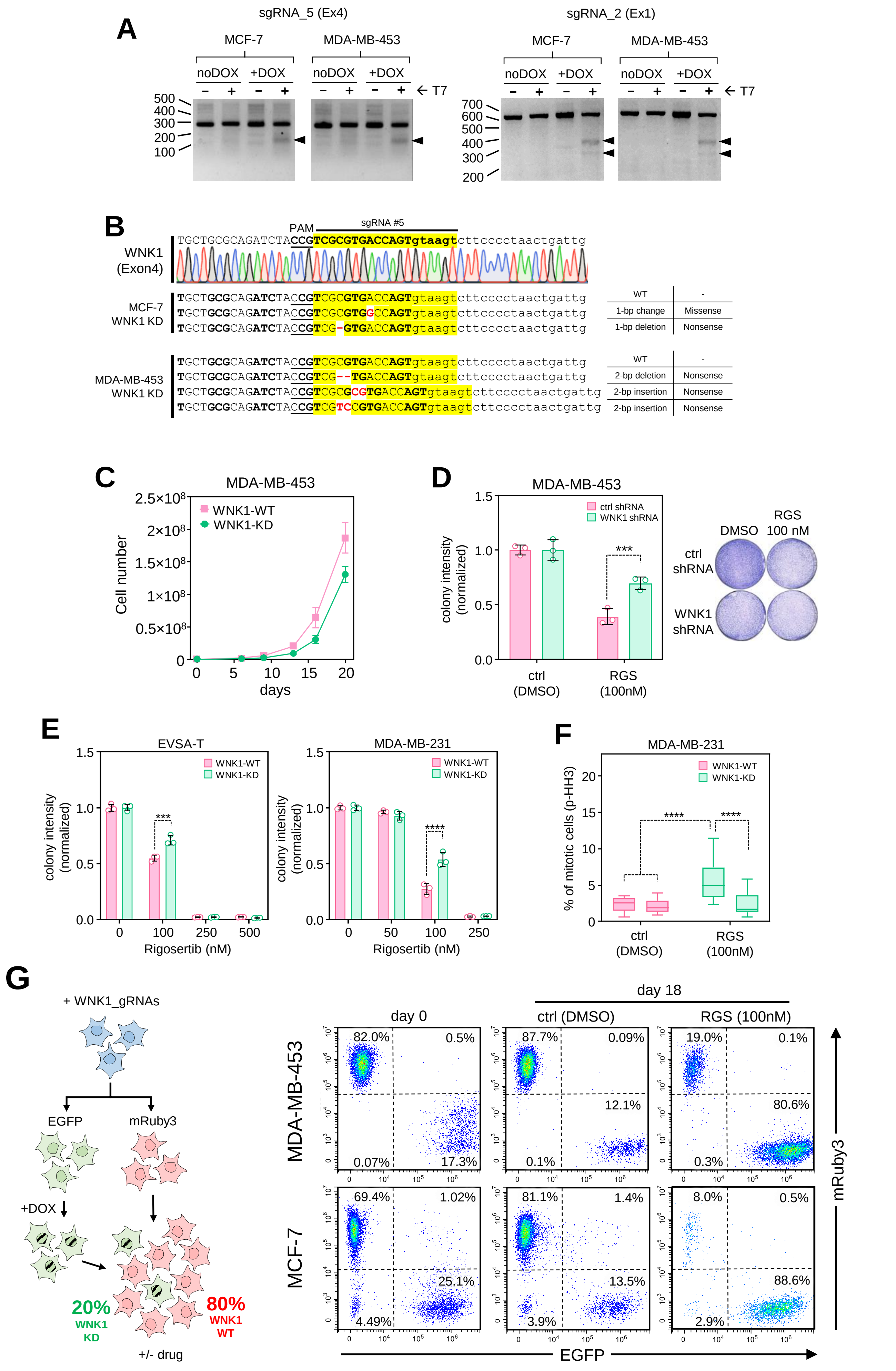
A
sgRNA_5 (Ex4)
sgRNA_2 (Ex1)
MCF-7
MDA-MB-453
noDOX
+DOX
T7
500
400
300
200
100
700
600
B
PAM
sgRNA #5
WNK1 (Exon4)
TGCTGCGCAGATCTACCGTCGCGTGACCAGTgtaagtcttcccctaactgattg
MCF-7 WNK1 KD
TGCTGCGCAGATCTACCGTCGCGTGACCAGTgtaagtcttcccctaactgattg
TGCTGCGCAGATCTACCGTCGCGTGGCCAGTgtaagtcttcccctaactgattg
TGCTGCGCAGATCTACCGTCG-GTGACCAGTgtaagtcttcccctaactgattg
WT
-
1-bp change
Missense
1-bp deletion
Nonsense
MDA-MB-453 WNK1 KD
TGCTGCGCAGATCTACCGTCGCGTGACCAGTgtaagtcttcccctaactgattg
TGCTGCGCAGATCTACCGTCG--TGACCAGTgtaagtcttcccctaactgattg
TGCTGCGCAGATCTACCGTCGCGCGTGACCAGTgtaagtcttcccctaactgattg
TGCTGCGCAGATCTACCGTCGTCCGTGACCAGTgtaagtcttcccctaactgattg
2-bp deletion
2-bp insertion
C
MDA-MB-453
Cell number
WNK1-WT
WNK1-KD
days
D
colony intensity (normalized)
ctrl shRNA
WNK1 shRNA
ctrl (DMSO)
RGS (100nM)
DMSO
RGS 100 nM
E
EVSA-T
MDA-MB-231
Rigosertib (nM)
F
% of mitotic cells (p-HH3)
G
+ WNK1_gRNAs
EGFP
mRuby3
+DOX
20% WNK1 KD
80% WNK1 WT
+/- drug
day 0
day 18
ctrl (DMSO)
RGS (100nM)
82.0%
0.5%
0.07%
17.3%
87.7%
0.09%
12.1%
0.1%
19.0%
0.1%
80.6%
0.3%
69.4%
1.02%
25.1%
4.49%
81.1%
1.4%
13.5%
3.9%
8.0%
0.5%
88.6%
2.9%
MCF-7
EGFP
mRuby3

Supplementary Figure 2

WNK1 depletion characterization and impact on rigosertib response:

(A) T7-endoculease assays of MCF-7 and MDA-MB-453 confirming the activity of Cas9 and WNK1 DNA edition at by sgRNA_5 (exon 4) and sgRNA_2 (exon 1).

(B) WNK1 exon 4 indels identification by DNA sequencing, in both MCF-7 and MDA-MB-453 cell lines.

(C) Growth curve of WNK1-WT and WNK-1KD MDA-MB-453 cells.

(D) Survival test for rigosertib by colony formation assay, of MDA-MB-453 cells expressing a WNK1 shRNA or a control shRNA. Two-way ANOVA test: $p<0.001$ (***)

(E) Survival test for rigosertib by colony formation assay, of WNK1-WT and WNK1-KD EVSA-T and MDA-MB-231 cells. Two-way ANOVA test: $p<0.001$ (***) and $p<0.0001$ (****)

(F) Mitotic index quantification by phospho-Ser10 histone H2 (p-HH3) staining in WNK1-WT and WNK1-KD MDA-MB-231 cells. Two-way ANOVA test: $p<0.0001$ (****)

(G) Schematic cartoon describing the cell competition assays (left panel) and the flow cytometry plots for EGFP and mRuby3 signal quantification (right panel).

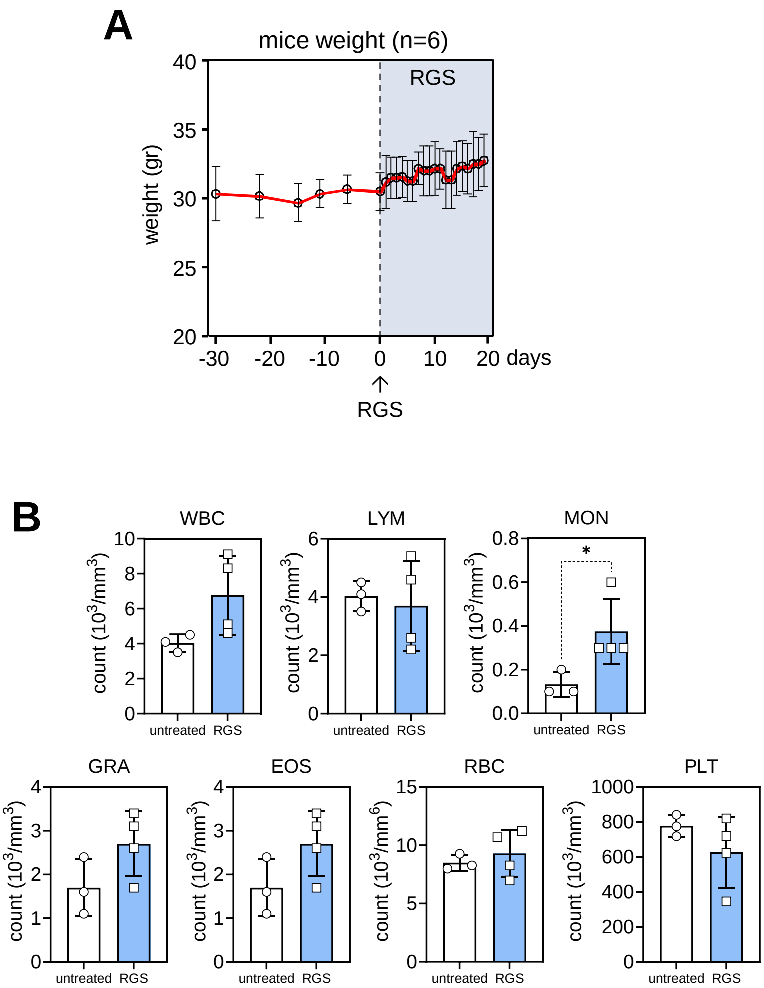
A
mice weight (n=6)
RGS
weight (gr)
40
35
30
25
20
-30
-20
-10
0
10
20 days
RGS
B
WBC
LYM
MON
GRA
EOS
RBC
PLT
count (10^3/mm^3)
count (10^6/mm^3)
untreated
RGS
*

Supplementary Figure 3

Analysis of rigosertib toxicity in xenograft mice.

(A) Animal weight measurements during the tumoral growth and the rigosertib treatment (blue area)

(B) Quantification of peripheral blood cell populations in untreated (white bars) and rigosertib treated mice (RGS - blue bars) at the end of the experimental setting. (WBC) White Blood Cells. (LYM) Lymphocytes. (MON) Monocytes. (GRA) Granulocytes. (EOS) Eosinophils. (RBC) ted Blood Cells. (PLT) Platelets. Two-way ANOVA test: $p<0.05$ (*).

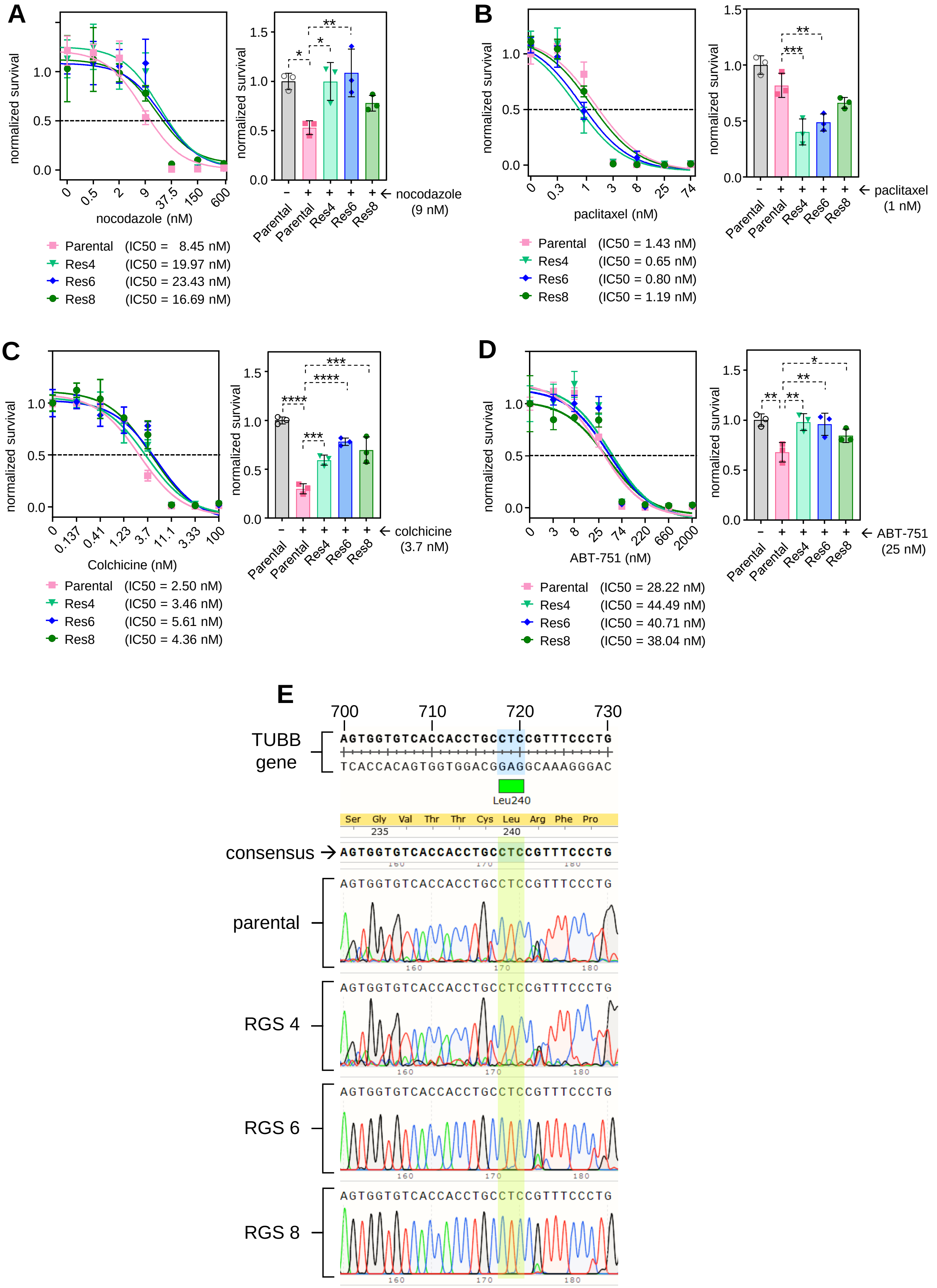
A
normalized survival
nocodazole (nM)
0
0.5
2
9
37.5
150
600
Parental (IC50 = 8.45 nM)
Res4 (IC50 = 19.97 nM)
Res6 (IC50 = 23.43 nM)
Res8 (IC50 = 16.69 nM)
← nocodazole (9 nM)
Parental
Parental
Res4
Res6
Res8
B
normalized survival
paclitaxel (nM)
0
0.3
1
3
8
25
74
Parental (IC50 = 1.43 nM)
Res4 (IC50 = 0.65 nM)
Res6 (IC50 = 0.80 nM)
Res8 (IC50 = 1.19 nM)
← paclitaxel (1 nM)
C
normalized survival
Colchicine (nM)
0
0.137
0.41
1.23
3.7
11.1
3.33
100
Parental (IC50 = 2.50 nM)
Res4 (IC50 = 3.46 nM)
Res6 (IC50 = 5.61 nM)
Res8 (IC50 = 4.36 nM)
← colchicine (3.7 nM)
D
normalized survival
ABT-751 (nM)
0
3
8
25
74
220
660
2000
Parental (IC50 = 28.22 nM)
Res4 (IC50 = 44.49 nM)
Res6 (IC50 = 40.71 nM)
Res8 (IC50 = 38.04 nM)
← ABT-751 (25 nM)
E
700
710
720
730
TUBB gene
AGTGGTGTCACCACCTGCCTCCGTTTCCCTG
TCACCACAGTGGTGGACGGAGGCAAAGGGAC
Leu240
Ser Gly Val Thr Thr Cys Leu Arg Phe Pro
235
240
consensus →
AGTGGTGTCACCACCTGCCTCCGTTTCCCTG
parental
AGTGGTGTCACCACCTGCCTCCGTTTCCCTG
RGS 4
AGTGGTGTCACCACCTGCCTCCGTTTCCCTG
RGS 6
AGTGGTGTCACCACCTGCCTCCGTTTCCCTG
RGS 8
AGTGGTGTCACCACCTGCCTCCGTTTCCCTG

Supplementary Figure 4

Response of isolated rigosertib resistant colonies to other microtubule-related drugs.

(A) Colony formation and IC50 calculation of parental MDA-MB-453 cell and rigosertib resistant cells, in the presence of nocodazole. Two-way ANOVA test: $p<0.05$ (*), $p<0.01$ (**)

(B) Colony formation and IC50 calculation of parental MDA-MB-453 cell and rigosertib resistant cells, in the presence of paclitaxel. Two-way ANOVA test: $p<0.01$ (**), $p<0.001$ (***)

(C) Colony formation and IC50 calculation, of parental MDA-MB-453 cell and rigosertib resistant cells, in the presence of colchicine. Two-way ANOVA test: $p<0.001$ (***), $p<0.0001$ (****)

(D) Colony formation and IC50 calculation of parental MDA-MB-453 cell and rigosertib resistant cells, in the presence of ABT-751. Two-way ANOVA test: $p<0.05$ (*), $p<0.01$ (**)

(E) DNA sequencing to test for the absence of the Leu240Phe mutation in the beta-tubulin gene (TUBB) in parental and rigosertib resistant isolated MDA-MB-453 cells.
